# Investigating the Influence of the Brain-Derived Neurotrophic Factor Val66Met Single Nucleotide Polymorphism on Familiarity and Recollection Event-Related Potentials

**DOI:** 10.1101/793356

**Authors:** Nicole S. McKay, David Moreau, Paul M. Corballis, Ian J. Kirk

## Abstract

The Val66Met single nucleotide polymorphism (SNP) has previously been reported to impact performance on recognition memory tasks. Whether the two subprocesses of recognition—familiarity and recollection—are differentially impacted by the Val66Met SNP remains unknown. Using event-related potentials (ERPs) recorded during a source memory task, we attempted to dissociate these two subprocesses. Behaviourally, we used participants’ scores on an item-recognition subtask as a measure of familiarity, and participants’ scores on a source-recognition subtask as a measure of recollection. Our findings reveal that Val/Val individuals outperform Met allele carriers on the item-but not the source-recognition task. Electrophysiologically, we were interested in the N400, an early frontal component previously linked to familiarity, and the late positive complex (LPC), a posterior component linked to recollection. We found evidence for Val/Val individuals having larger amplitudes of the LPC compared to Met allele carriers, and evidence for no difference in the N400 amplitudes of these groups. Based on the lack of dissociation between familiarity- and recollection-specific ERPs at the LPC time window, we argue that our behavioural and ERP results might reflect better item-recognition for Val/Val individuals compared to Met allele carriers. We further suggest that both these results reflect differences related to familiarity, rather than recollection.

## Introduction

Dual-process accounts of recognition memory propose that recognition of prior experience can be based on two types of information processing; familiarity and recollection (see Yonelinas et al., 2007, for a review). Familiarity is an automatic process that gives rise to a sense of whether or not an object has been previously experienced, independent of any contextual information. In contrast, recollection is a slower, more effortful process that involves the recall of additional episodic information. These two subprocesses of recognition have been behaviourally and structurally dissociated in previous research (Aggleton & Brown, 1999; Aggleton, Dumont, & Warburton, 2011; Brown & Xiang, 1998; Eichenbaum, Yonelinas, & Ranganath, 2007; Henson, Rugg, Shallice, Josephs, & Dolan, 1999; Hintzman, Caulton, & Levitin, 1998; Mitchell & Johnson, 2009; Wixted, 2007; Yonelinas, 1994; Yonelinas, Kroll, Dobbins, Lazzara, & Knight, 1999; Yonelinas, Otten, Shaw, & Rugg, 2005). A consistent behavioural distinction between these two subprocesses is that familiarity-based recognition is much faster, and can reflect a wide range of confidence levels, while recollection requires more processing time, and is only associated with high levels of confidence (Yonelinas, 2002). Structurally, familiarity has been demonstrated to be especially reliant on the perirhinal cortex, while recollection is critically dependent upon the hippocampus (see Aggleton and Brown, 1999, for a review).

In addition to behavioural and structural distinctions, many studies have reported that familiarity and recollection are characterised by unique electrophysiological correlates (Addante, Ranganath, & Yonelinas, 2012; Cansino & Trejo-Morales, 2008; Curran, 2000; Friedman & Johnson, 2000; Hoppstädter, Baeuchl, Diener, Flor, & Meyer, 2015; Leynes, Landau, Walker, & Addante, 2005; Rugg & Curran, 2007; Wilding & Rugg, 1996; Yu & Rugg, 2010). Event-related potentials (ERPs) recorded during recognition memory tasks are one method used to investigate functional dissociations of these two subprocesses of recognition. This method is particularly useful as familiarity and recollection have unique temporal properties; both these processes are thought to emerge in the order of tens of milliseconds after stimulus onset. More specifically, at retrieval familiarity has been linked to a frontal, negative, ERP component that occurs approximately 400ms post-stimulus, known as the N400 (Friedman & Johnson, 2000; Gruber & Otten, 2010; Rugg & Curran, 2007; Voss & Paller, 2007). In contrast, recollection has been linked to a late positive complex (LPC) that arises 500-800ms post-stimulus at posterior-parietal electrodes, particularly over the left hemisphere (Addante et al., 2012; Curran, 2000). These two components have been reported across a large number of recognition experiments using a variety of measures, such as confidence ratings, to discriminate familiarity from recollection (Addante et al., 2012; Düzel, Yonelinas, Mangun, Heinze, & Tulving, 1997; Wilding & Rugg, 1996; Woodruff, Hayama, & Rugg, 2006; Yu & Rugg, 2010).

One paradigm often used to dissociate familiarity and recollection within a single trial is the source memory task (Addante et al., 2012; Johnson, Hashtroudi, & Lindsay, 1993). This task requires participants to not only recognise previously presented items but also to retrieve associated episodic information about the encoding period of that item. Specifically, familiarity is important for the initial recognition of a studied item regardless of whether the encoding period is explicitly remembered (item-recognition). In contrast, recollection is required for the recognition of episodic details from the encoding period for each item (source-recognition). By comparing performance on trials in which items were correctly identified as “old” with correct source-recognition, to trials in which items were correctly identified as “old” without correct source-recognition, we can get a measure that indexes recollection. These assumptions have been supported by a number of ERP studies that suggest that performance on an item-recognition judgment is associated with modulations of the N400, while source-recognition judgment is associated with the LPC (Addante et al., 2012; Eichenbaum et al., 2007; Mollison & Curran, 2012; Woodruff et al., 2006). More specifically, these studies have found that the N400 occurs for all correctly recognised items, while the LPC is largest when items are correctly remembered alongside details of their encoding context.

Given the above-mentioned studies, it is interesting to consider how individual differences in recognition memory might relate to the familiarity and recollection subprocesses of the source memory task. One source of performance differences could be common genetic variations within the population (Payton, 2006). For example, a single nucleotide polymorphism (SNP) known as the Val66Met SNP (rs6265), occurs within the gene coding for brain-derived neurotrophic factor (BDNF) and has been previously related to reduced accuracy on a large number of memory tasks (Dempster et al., 2005; Egan et al., 2003; Hariri et al., 2003; Kennedy et al., 2015; Lamb, Thompson, McKay, Waldie, & Kirk, 2015). Several recent studies have reported that carriers of this SNP perform worse than non-carriers on recognition memory tasks (Komulainen et al., 2008; Spriggs et al., 2019). The Val66Met SNP is a non-synonymous SNP that results in a switch from a valine (Val) to methionine (Met) amino acid. This new amino acid impacts the trafficking of the resulting BDNF protein and its activity-dependent secretion (Chen et al., 2004). These biological changes are thought to influence memory formation by impacting context-specific synaptic plasticity (Schofield et al., 2009; Spriggs et al., 2019), an important mechanism underpinning learning and memory.

The influence of the Val66Met SNP on brain electrophysiology has been previously measured using both oscillatory EEG activity and ERPs recorded during a variety of cognitive tasks (Beste et al., 2010; Gajewski, Hengstler, Golka, Falkenstein, & Beste, 2012; Gatt et al., 2008; Soltész et al., 2014). For example, carriers of the Met allele are reported to have decreased oscillatory activity and synchrony during error processing (Soltész et al., 2014), and weaker error-specific phase locking (Beste et al., 2010). Other studies have reported Met allele carriers have increased latency and reduced amplitude for the P300, an ERP with generators that include the hippocampus (Getzmann, Gajewski, Hengstler, Falkenstein, & Beste, 2013; Schofield et al., 2009; Soltész et al., 2014). Furthermore, serum levels of BDNF have been positively correlated with the power of gamma oscillations (Hiramoto et al., 2017). Despite the abundance of EEG-derived research showing a relationship between BDNF and electrophysiological measures, little is known about how this SNP influences the electrophysiology of the brain during recognition.

We sought to test whether BDNF genotype differentially impacted familiarity- and recollection-based recognition judgments. We used a source memory paradigm to differentiate between these two subprocesses both on the behavioural and electrophysiological levels. We hypothesised that individuals with two copies of the valine allele (Val/Val individuals) would perform with increased accuracy on our overall recognition memory task, and specifically the two subscores of item-recognition and recognition of source context, compared to Met allele carriers. In particular, we predicted that the accuracy difference between our genotype groups would be greatest for the source-recognition subtask, a recollection-based judgment, because of the association between recollection processing and the hippocampus (Aggleton & Brown, 1999). Previously, context-dependent BDNF secretion, which is important for memory, had been shown to be highest in this region (Chen et al., 2004). We further hypothesised that there would be differences in the average ERP amplitudes of Val/Val and Met carrying participants calculated across the N400 and LPC time windows. We predicted that Val/Val individuals would have larger amplitudes, and a greater magnitude of the old/new effect, compared to Met allele carriers. This is in line with previous research that has found increased amplitudes on this task to be correlated with increased accuracy. Specifically, we predicted that the differences in these ERP amplitudes might be greatest for the LPC, the ERP component linked to recollection processing, compared to the N400, the component related to familiarity.

## Methods

### Participants

Seventy-seven healthy adults, aged 18-33 (*M* = 23.3, *SE* = 0.50), volunteered to complete this study. Nine participants were excluded from the study due to insufficient task performance (3) or technical issues with preprocessing their EEG data (6). Demographic information for the remaining 69 participants included in this study can be viewed in Table 1. All participants were recruited through the University of Auckland and reported no history of neurological disorder. Genotyped participants were divided into two groups, those with the Val/Val genotype, and those with at least one copy of the Met allele. A subset of sixty-one of the participants that completed the EEG task also participated in an MRI, which has been previously reported (McKay, Moreau, Henare, & Kirk, 2019). All participants gave informed consent, and this study was approved by the University of Auckland Human Participants Ethics Committee.

**Table 1.** General participant information for participants included in the EEG analyses.

|  | Whole sample | Val/Val | Met Carriers | *p*-values |
| --- | --- | --- | --- | --- |
| N | 68 | 37 | 31 |  |
| Mean Age (SE) | 23.4 (0.50) | 24.1 (0.67) | 22.4 (0.72) | .09 |
| Females (Males) | 45 (23) | 22 (15) | 23 (8) | .20 |
| Right (Left) Handed | 59 (9) | 32 (5) | 27 (4) | .94 |

Note: *p*-values are derived from testing differences between the two genotype groups using an independent-samples *t*-test for age; and chi-squared analyses for sex, handedness, ethnicity.

### Genotyping

Saliva samples were collected from all participants using Oragene-DNA Self Collection kits and stored at room temperature until analysis could take place. Prior to collection, participants were asked not to eat or drink for at least 30 minutes. DNA was extracted from the sample and was resuspended in Tris-EDTA buffer and quantified using a Nanodrop ND-1000 1-position spectrophotometer (Thermo Scientific). Amplification was conducted on the 113bp fragment that coincides with the Val66Met SNP within the BDNF gene. The primers used for this amplification were *BDNF-F 5’-GAG GCT TGC CAT CAT TGG CT-3’ and BDNF-R 5’-CGT GTA CAA GTC TGC GTC CT-3’*. Polymerase chain reaction (PCR) was then applied to the samples, and enzyme digestion was used to cut the samples into the relevant sections. Val fragments of the DNA samples result in two sections of differing lengths, one of 78bps and the other of 35bps, while Met fragments resulted in one section of 113bps in length. DNA was then visualised under ultraviolet light and participants were classified as either Val/Val or Met carriers (those with either Val/Met or Met/Met genotypes). Based on the resulting DNA visualisation, our participants were classified as either Val/Val or Met carriers resulting in 37 Val/Vals and 31 Met carriers.

### Source Memory Task

We used a modified version of the source memory task outlined in Addante et al. (2012). This task requires participants to learn items as well as make an associated judgment, and allows for two types of recognition to be assessed (see Figure 1). Item recognition thought to index familiarity, and source-recognition thought to index recollection. During the study phase of the experiment, participants were presented with 300 objects and were asked to make an associated judgment for each object. On 150 of these trials, participants were asked to decide whether the object presented to them was man-made, and on the remaining 150 trials, participants were asked to decide whether the object presented could fit in a box (size specified by the researcher). Objects were presented to participants in six blocks of 50 stimuli, with breaks of at least 30 seconds between each block. Prior to each stimulus presentation, a fixation cross was presented. The encoding probe (man-made or box) then appeared on the screen for 400 ms. Following this, participants were presented with the object for 1500 ms, and following the trial, participants were asked to respond with a “yes” (press 1) or “no” (press 2) to the cued judgment for that trial. After the study phase, participants were given a 30-minute break with a distraction task (sudoku), and light refreshments were provided. During the retrieval phase, the 300 objects presented in the study phase were mixed with a further 100 novel objects. These 400 objects were presented to the participants in eight blocks of 50. Before each item presentation, a fixation cross was presented for 750 ms. The object was then presented for 1500 ms. At the end of the trial, participants were asked to decide whether the object presented was from the previous study phase, “old” (press 1), or was a new picture, “new” (press 2). If participants responded that the object was old, a second response screen was presented that asked with which of the two judgments the object was paired within the study phase, “man-made” (press 1), or “box” (press 2). All trials were preceded by a blank screen that lasted between 750 and 1250 ms, to minimise any expectation bias influencing responses.

**Figure 1.**
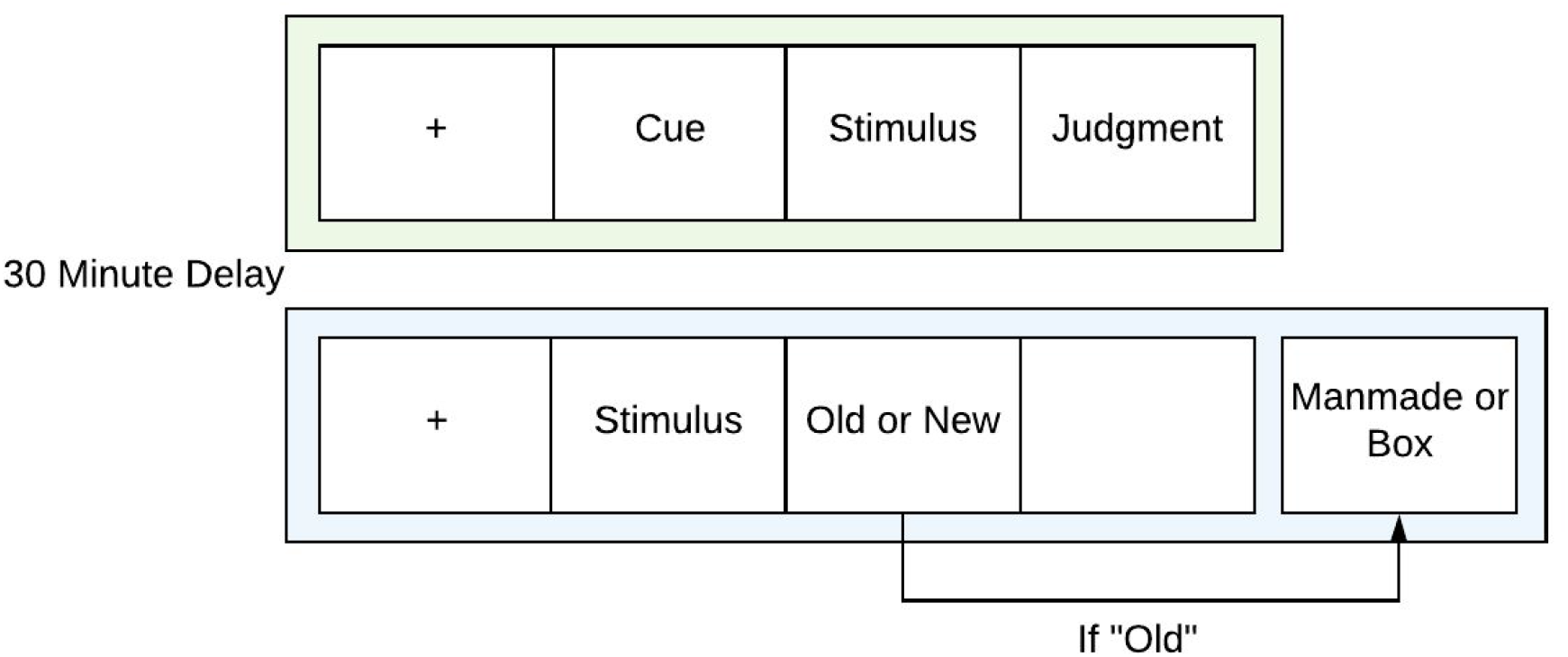
Source recognition memory paradigm. A schematic depiction of the paradigm used for the experiment (adapted from Addante et al., 2012). Participants were shown 300 items in the encoding phase (green) and asked to make an associated judgment for each. After a 30 minute delay, participants then completed the retrieval phase of the experiment (blue).

All key presses were made by participants on a standard keyboard number pad, and participants were given unlimited time to respond within each trial. All testing was conducted in a dimly lit, electrically shielded room. Stimuli were presented on an LCD monitor with screen dimensions of 47.7 x 26.8cm, and participants were positioned 57cm away from the screen to control for visual angle. E-prime (v 2.0.8.74) was used to control stimulus presentation and synchronisation with the EEG recording and participants’ responses.

### Stimuli

Picture stimuli were selected from the Bank of Standardized Stimuli (BOSS) (Brodeur, Dionne-Dostie, Montreuil, & Lepage, 2010). This database consists of 930 photos of everyday objects that have been normalised for name, category, familiarity, visual complexity, object agreement, viewpoint agreement, and manipulability (Brodeur, Guérard, & Bouras, 2014). All images are edited so that luminance and colour are equal across the images. Of these 930 photo stimuli, a subset of 500 images was randomly chosen as stimuli for this study. Two images were removed after pilot participants’ feedback that the object was not known in a New Zealand context. Each participant was exposed to a random selection of 400 of these images, 300 in the encoding phase, and a further 100 were introduced in the recognition phase.

### Data Acquisition

EEG activity was recorded using an Electrical Geodesics Inc. (EGI) system (EGI Inc., Eugene, Oregon, USA). A high-density 128-channel Ag/AgCl HydroCel Geodesic Sensor Net was used for the recording, and each participant’s head circumference was measured to ensure an appropriate electrode net was applied. The signal was recorded at 1000Hz with an online reference electrode located at the vertex (Cz) and a ground electrode (COM) located between the vertex and the Pz electrodes. EEG activity was amplified with an EGI net amps 400 amplifier with 24-bit A/D conversion, an input impedance of ≥ 200 MΩ, and a 4kHz antialiasing filter (500Hz low pass cut off). Electrode impedance levels were kept below 40kΩ, and EGI’s implementation of the driven common technique (Common Active Noise Sensing and Limiting) was used throughout.

### Data Processing

All EEG data were processed within EEGLAB (Delorme & Makeig, 2004), an analysis toolbox that works within the Matlab environment (2016a, Mathworks Inc.). EEG data were processed twice, once in a manner specialised for independent component analysis (ICA), and once in a manner specialised for ERP analysis. This was necessary as optimal preprocessing for ICA denoising requires a 1Hz high-pass filter to be applied to the data, which is known to distort late ERP components such as the N400 and LPC. After our data were analysed using ICA, the resulting weights were transferred to data resulting from a more conventional ERP preprocessing pipeline (Luck, 2014). This allowed us to use the components from the ICA to remove eye and muscle artefacts, without having to apply unsuitable filters to our data (see: https://sccn.ucsd.edu/wiki/Makoto's_preprocessing_pipeline for pipeline overview). Specific processing parameters for each stream are outlined below.

#### ICA preprocessing

EEG data were first downsampled to 250Hz. A 1Hz high-pass filter was applied to remove baseline drift (Winkler, Debener, Müller, & Tangermann, 2015). Channel information was imported, and channels with excessive noise were identified and removed. In order to accomplish this, we calculated and removed any channel that had spectral power between 0-35Hz and was more than 15 standard deviations from the mean of our other channels. Then we calculated peak-to-peak differences for each channel in windows of 5000 sampling points, across the entire length of recorded data. If any electrode’s peak-to-peak difference was calculated as more than 10 standard deviations from the mean, more than 5 times, it was removed. For any channel that was removed, a spherical interpolation of the surrounding channels was introduced in its place. Data was then re-referenced to the average of all electrodes, and line noise was removed using the CleanLine plugin (Mullen, 2012). ICA was then performed with the PCA option selected to reduce data dimensions due to previously interpolated channels, and also so we would only examine the top 30 contributing components.

#### ERP preprocessing

EEG data was kept at the original 1000Hz sampling rate. A 0.1-30Hz Butterworth bandpass filter was applied using the ERPLAB plugin for EEGLAB (Lopez-Calderon & Luck, 2014). Channel location information was imported, and data were epoched 200ms pre-stimulus to 1500ms post-stimulus. Channels identified as containing bad data were removed and interpolated following the same criteria as stated above. The ICA variables calculated via the ICA preprocessing steps were transferred to this data, and ICA components associated with artefacts were identified by the ADJUST toolbox (Mognon, Jovicich, Bruzzone, & Buiatti, 2011). All ICA components identified by this toolbox were manually inspected, and only those that fell within the 10 largest contributing components, were rejected, resulting in an average of 3.6 ICA components being rejected per person. Epochs that contained amplitudes that exceed +/- 100μV were also rejected. Data were then re-referenced to the average of all electrodes. Data were then epoched into the following condition types: correctly identified new items (Correct Rejections), correctly identified old items with correctly recognised source judgment (Hit-Hit), correctly identified old items with incorrectly recognised source judgment (Hit-Miss), and incorrectly identified old items (Missed). We were unable to epoch our data reliably into a condition for incorrectly identified new items (False Alarms) because many participants had a small number of False Alarm responses. Importantly, the conditions we were able to epoch enabled us to make comparisons between both correctly identified old and new items, as well as between correctly identified old items with correct source judgment, and correctly identified old items with incorrect source judgment. Participants with less than 10 trials for any condition were removed, leading to five individuals being dropped from our analyses. The remaining participants had an average of 81 trials per condition.

We used *a priori* defined time windows and electrode sites of interest based on previous research dissociating familiarity and recollection using ERPs (Addante et al., 2012). ERPs were derived by taking the average of a cluster of electrodes from each of the regions of interest (Figure 2). Familiarity has been previously localised to frontal, central regions, so a cluster of seven electrodes centred on electrode 6 was used to produce familiarity-related ERPs. This cluster corresponds to a region that would fall between the locations of the Cz and Fz electrodes of the International 10-10 system. In contrast, recollection has been previously localised to posterior, left-lateralised regions, so a cluster of seven electrodes centred on electrode 60 was used to produce recollection-related ERPs. This cluster corresponds to a region that would fall approximately between the locations of the Pz and P3 electrodes of the International 10-10 system. Additionally, each subprocess of recognition has been previously linked to specific time windows of interest, therefore, old/new effects were examined for two time windows of 300-500ms post-stimulus and 500-800ms post-stimulus. Analyses examining differences between familiarity (Hit-Miss) and recollection (Hit-Hit) ERPs were conducted on the late 500-800ms time window.

**Figure 2.**
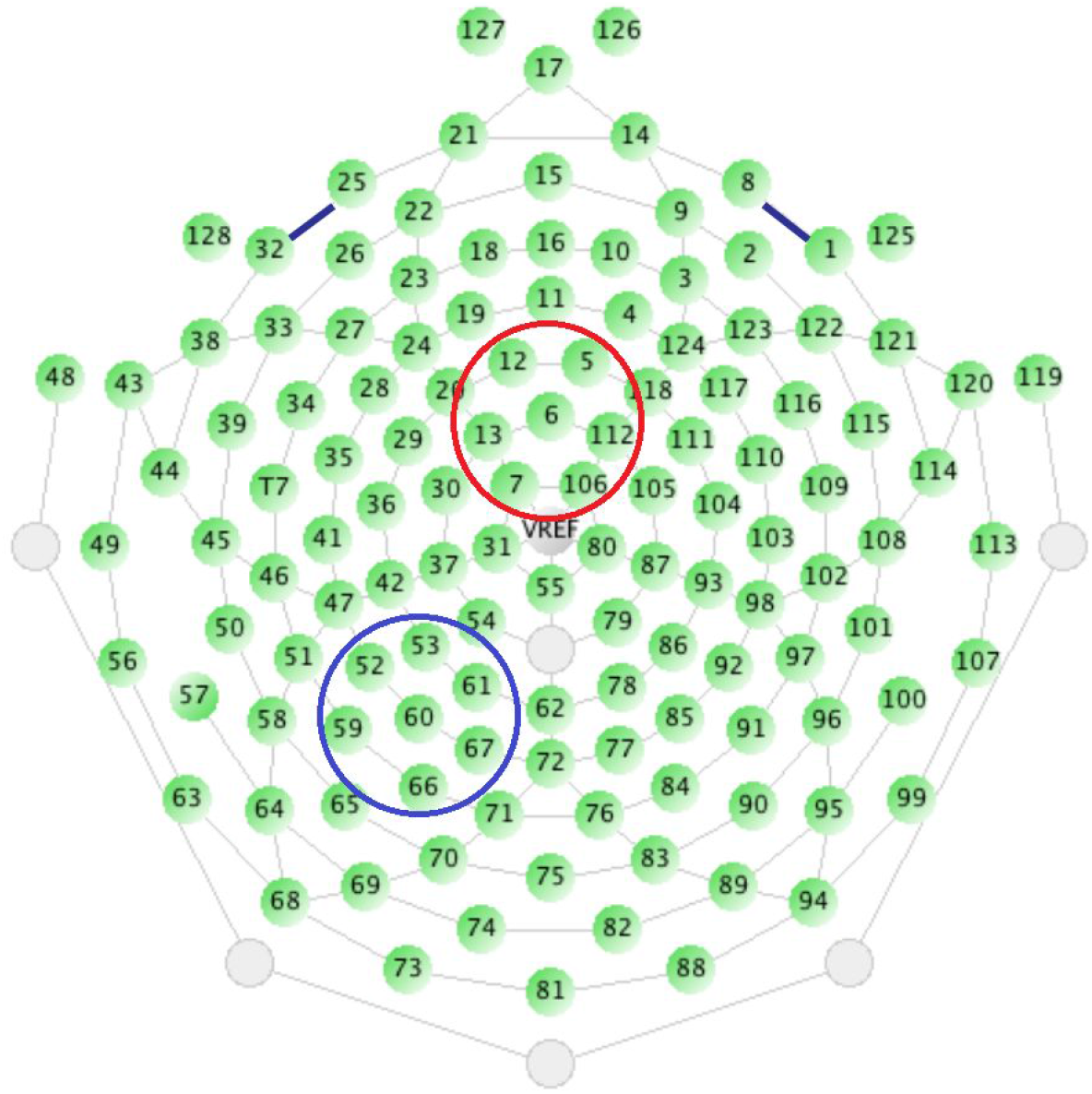
Specific electrode locations for each time window of interest. Electrodes mapped in red are the cluster of seven electrodes used for the N400 analyses, while electrodes mapped in blue are the cluster of seven electrodes used for the LPC analyses.

### Analyses

Given the experimental setup, signal detection measures were derived from the resulting behavioural data on both subtasks. Hit and Miss rates were calculated as the proportion of the old items that were correctly and incorrectly recognised. Correct rejections (CR) and False Alarms (FA) were calculated as the proportion of new items that were correctly identified as new, and the proportion of new items that were incorrectly recognised as old. In order to examine whether there was an impact on an individual’s ability to discriminate between old and new items, we derived a measure of sensitivity (*Pr*) by subtracting False Alarms from the Hit rate on the item-recognition task. Our sensitivity measure allowed us to ensure any potential results for differences in Hit rate were not an artefact of participants just responding “old” for all items, and also allowed us to take participant errors into account when considering how well they performed on the task. Finally, we derived a measure of response bias (*Br*) using the following equation:

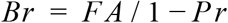

The above measures were calculated for the item-recognition subtask and therefore were independent of whether a participant also scored correctly on the source-recognition task. In order to compare the relationship between accuracy on the item-recognition and source-recognition tasks, we first manipulated our item-recognition scores so that both the item-recognition and source-recognition tasks had a chance accuracy rate of 50%. Item recognition accuracy was calculated as the sum of Hits and Correct Rejections, divided by the total number of trials. Source recognition accuracy was calculated calculating the proportion of items that were correctly recognised as old with correct source-recognition, divided by the number of correctly recognised old items.

All statistical analyses were conducted in JASP (JASP Team, 2016), and subsequent plots were made using R (R Core Team, 2019). We used Bayesian statistics for all our analyses, however, the frequentist equivalents are provided in the Supplementary Material to allow the reader to compare results across frameworks. To analyse the behavioural data we used six separate Bayesian independent-samples *t*-tests with the default Cauchy scale (*r* = 0.707; Morey & Rouder, 2018), one for each unique behavioural measure; Hits, CR, *Pr, Br*, Item Accuracy, Source Accuracy. The additional behavioural measures were not subjected to further tests as they are complementary measures to the ones included. To analyse the ERP results we first examined differences between the amplitudes for ERPs evoked by correctly recognised old (Hits) vs. new (Correct Rejections) items for each time window of interest, using two separate Bayesian paired-samples *t*-tests with the default Cauchy scale (*r* = 0.707). To specifically compare recollection and familiarity, we ran a further Bayesian paired-samples *t*-test with the default Cauchy scale (*r* = 0.707) on peak amplitudes of ERPs evoked by items correctly recognised with source-recognition (Hit-Hit), and items correctly recognised without source-recognition (Hit-Miss). Finally, to examine how Val66Met genotype influences these effects, we performed three separate Bayesian independent-samples *t*-tests with the default Cauchy scale (*r* = 0.707). These *t*-tests were performed on difference scores computed across the time windows of interest. The first two looked at the difference between old and new ERPs for the N400 and LPC time windows, and the third looked at the difference between familiarity (Hit-Miss) and recollection (Hit-Hit) ERPs across the LPC time window.

## Results

### Behavioural Analyses

We ran six separate Bayesian independent-samples *t*-tests to quantify the behavioural evidence for Val66Met group differences in recognition memory. Specifically, we were interested in evaluating the relative support for the null hypothesis that there is no difference between the two genotypes for our accuracy measures, and also the hypothesis that Val/Val participants scored higher on accuracy measures than Met carriers. Our response bias measure was subjected to a two-tailed independent-samples *t*-test as we had no directional hypothesis *a priori*.

For both the Hit and Correct Rejection rates we found anecdotal evidence in favor of Val/Val participants having higher accuracy compared to Met allele carriers (Hit: BF_10_ = 1.2, *ε* = 8.0×10^-3^%, *M* = 0.77, *SD* = 0.10 and *M* = 0.73, *SD* = 0.13, respectively; Correct rejection: BF_10_ = 1.4, *ε* = 6.2×10^-5^%, *M* = 0.93, *SD* = 0.04 and *M* = 0.90, *SD* = 0.07, respectively). This can be interpreted as weak evidence that Val/Val individuals were better at recognising old items, as well as better at identifying new items, compared to Met allele carriers. In line with this, for our measure of discriminability, *Pr*, we found moderate support for Val/Val individuals being better able to discriminate old items from new items compared to Met allele carriers (BF_10_ = 4.4, *ε* = 4.9×10^-4^%, *M* = 0.70, *SD* = 0.10 and *M* = 0.63, *SD* = 0.14, respectively). Our measure of item-recognition accuracy also supports these results. We found anecdotal evidence for Val/Val individuals having greater accuracy on the item-recognition task (BF_10_ = 1.9, *ε* = 5.0×10^-3^%, *M* = 0.81, *SD* = 0.07 and *M* = 0.77, *SD* = 0.10, respectively). In contrast, for our source-recognition accuracy measure we find anecdotal evidence to support the null hypothesis that there is no difference in score for Val/Val compared to Met allele carriers (BF_01_ = 1.5, *ε* = 5.0×10^-3^%, *M* = 0.78, *SD* = 0.07 and *M* = 0.76, *SD* = 0.07, respectively). Finally, for the measure of response bias, *Br*, we found moderate support for the null hypothesis that there was no difference in response bias for each of the two genotype groups (BF_01_ = 3.9, *ε* = 6.0×10^-3^%, Val/Val: *M* = 0.26, *SD* = 0.16 and Met: *M* = 0.27, *SD* = 0.16). Distributions for the scores of each group for each measure are shown in Figure 3. Posterior and prior distributions, as well as sequential analyses and robustness checks for these tests, are available in the Supplementary Material (Figures SM.1:3).

**Figure 3.**
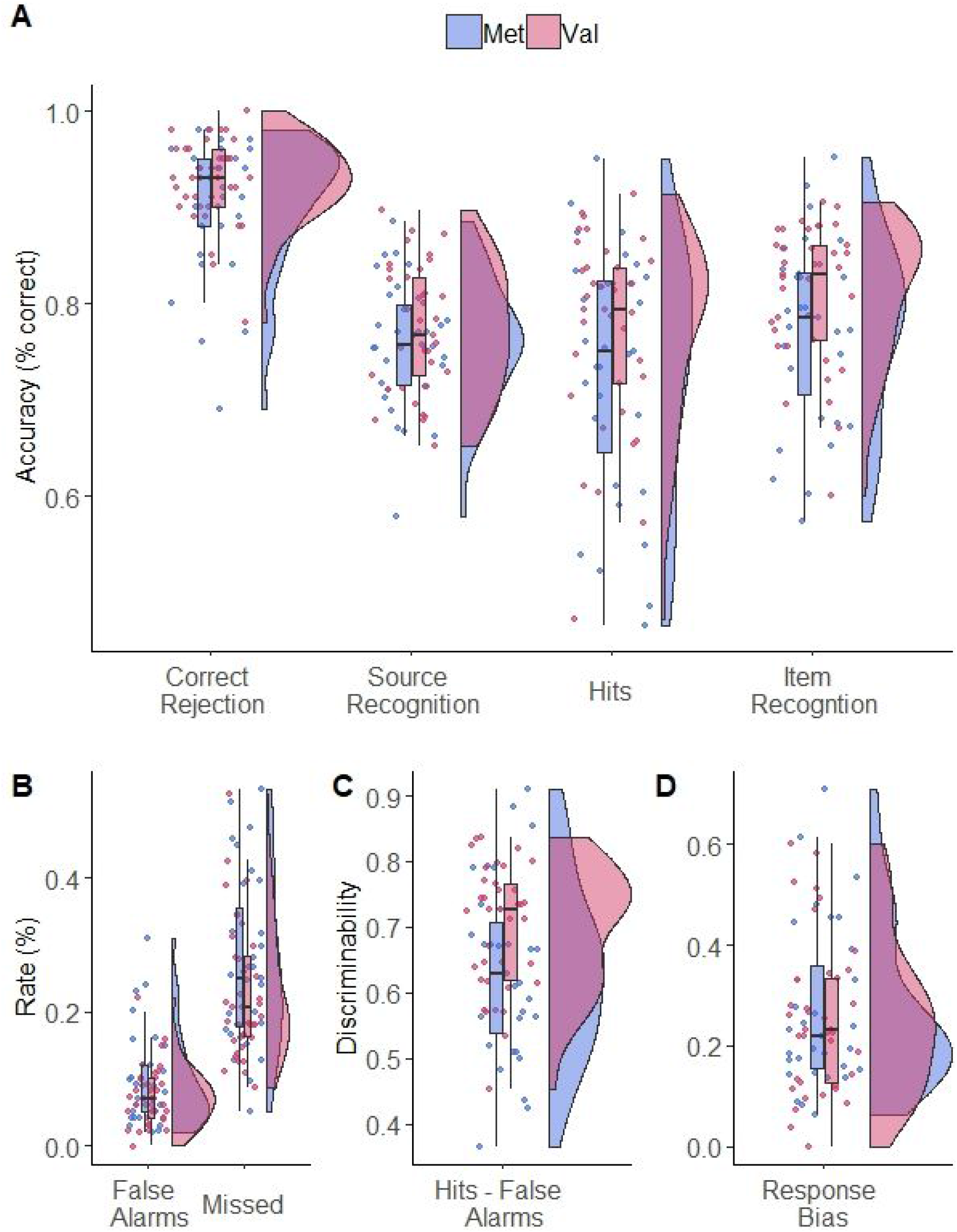
Distributions for each of the performance measures. **A:** Distributions of the accuracy scores across the source memory task. Correct rejections measure the percentage of trials with new items that participants correctly identified as new. Source accuracy measures the accuracy of the recollection specific subtask, while Item Accuracy measures the accuracy of the familiarity specific subtask. Finally, Hits are measuring the raw Hit rate for the overall source memory task. **B:** Distributions of the error scores across the source memory task. False alarms are incorrectly recognised new items, and Missed are items that were incorrectly identified as new. **C:** This panel shows distributions of discriminability, a measure of sensitivity calculated by subtracting False Alarms from the overall Hit rate. **D:** Distributions of response bias for each group. In all cases, Met carriers are depicted in blue, and Val/Val individuals are depicted in red.

### N400 Component

In order to test whether we replicated the early old/new effect reported in previous research, a Bayesian paired-samples *t*-test was conducted on the average amplitude of ERPs across the N400 time window in response to correctly identified old items, and correctly identified new items (Figure 4). We found strong evidence for a difference between the amplitudes of ERPs elicited in response to correctly identified old items compared to correctly identified new items (BF_10_ = 2.3×10^10^, *ε* = 6.0×10^-17^%, *M* =-1.7, *SD* = 1.7 and *M* = −2.3, *SD* = 1.8, respectively).

**Figure 4.**
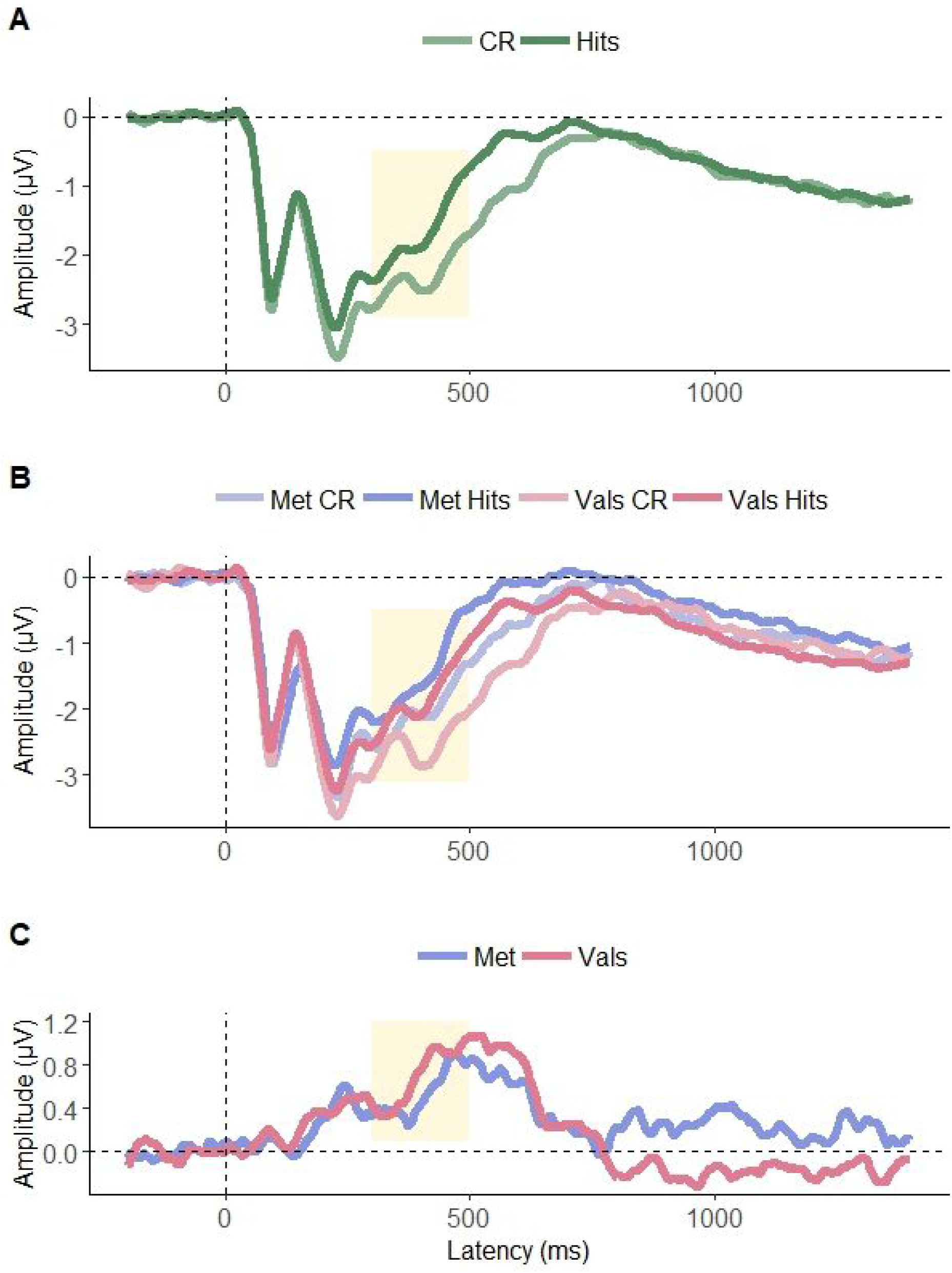
ERPs illustrating the electrophysiological responses to each of the correctly identified Old (Hits), and New (CR) items. **A:** ERPs for the overall group showing replication of previously reported old/new effects at frontal electrode sites. **B:** ERPs split by genotype demonstrating how each of Val/Val and Met allele individuals respond to different trial types. **C:** Difference waves calculated between ERPs elicited to old and ERPs elicited to new items. These are recorded at a cluster of electrodes centred around electrode 6, a central, frontal electrode. Time window of interest for the N400 component is highlighted in yellow (250-500ms).

#### Genetic impact on N400

To determine whether the Val66Met SNP impacted the N400 component of the ERPs gathered, we first determined the magnitude of the difference between the ERP evoked in response to correctly identified old items, and the ERP evoked in response to correctly identified new items. The average of this difference value across the N400 time window was then subjected to a Bayesian independent-samples *t*-test to examine whether the size of the old/new effect was different for the two Val66Met groups. We found anecdotal evidence for the null hypothesis that there is no difference in the magnitude of the old/new effect between Val and Met allele individuals (BF_01_= 1.4, *ε* = 3.0×10^-3^%, *M* =0.70, *SD* = 0.54 and *M* = 0.54, *SD* = 0.62, respectively).

Distributions for the scores of each group for each measure are shown in Figure 7. Posterior and prior distributions, as well as sequential analyses and robustness checks for these tests, are available in the Supplementary Material (Figure SM2).

### Late Positive Complex

In order to test whether we replicated the late old/new effect reported in previous research, we conducted a Bayesian paired-samples *t*-test on the average amplitude of ERPs across the LPC time window in response to correctly identified old items, and correctly identified new items (see Figure 5). We found moderate evidence for the absence of a difference between the amplitudes of ERPs elicited in response to correctly identified old items compared to correctly identified new items (BF_01_ = 5.4, *ε* = 3.7×10^-5^%, *M* = 3.1, *SD* = 2.1 and *M* = 3.0, *SD* = 2.2, respectively). We further aimed to examine whether we replicated the findings of Addante et al. (2012), that showed a dissociation in the LPC time window between correctly identified old items with remembered source judgment (Hit-Hit), and those where the associated judgment was not remembered (Hit-Miss). Therefore, a second Bayesian paired-samples *t*-test was conducted on the average amplitudes for each of these conditions across the LPC time window. We found further support for the null hypothesis of no difference in the mean amplitude for our conditions for the LPC time window (BF_01_ = 3.5, *ε* = 2.4×10^-5^%, Hit-Hit: *M* = 3.1, *SD* = 2.1 and Hit-Miss: *M* = 2.9, *SD* = 2.4, respectively). Therefore, we did not replicate the effect described in the Addante et al. (2012) paper.

**Figure 5.**
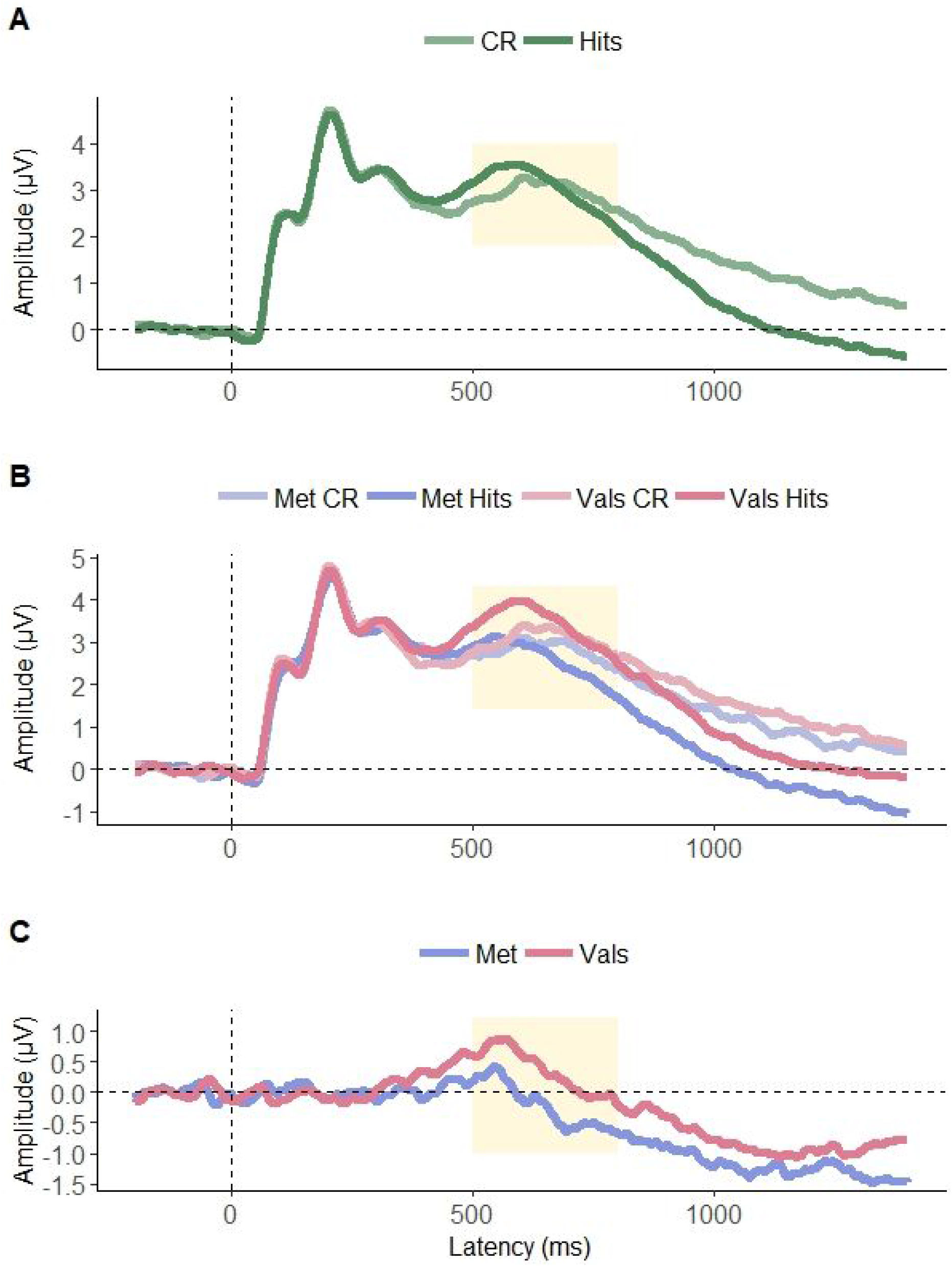
ERPs illustrating the electrophysiological responses to each of the correctly identified Old (Hits), and New (CR) items at the LPC electrode cluster. **A:** ERPs for the overall group showing replication of previously reported old/new effects at posterior electrode sites. **B:** ERPs split by genotype demonstrating how each of Val/Val and Met allele individuals respond to different trial types. **C:** Difference waves calculated between ERPs elicited to old and ERPs elicited to new items. These are recorded at a cluster of electrodes centred around electrode 60, a left-ward, posterior, electrode. Time window of interest for the LPC component is highlighted in yellow (500-800ms).

#### Genetic impact on LPC

In order to examine the impact of Val66Met genotype on recognition ERPs across the LPC time window, we first calculated difference scores across the time window of interest between ERP amplitudes elicited as a response to correctly identified old items and correctly identified new items. Using a Bayesian independent-samples *t*-test, we found anecdotal evidence for a difference in the magnitude of the old/new effect across the LPC time window for Val/Val individuals compared to Met allele carriers (BF_10_= 5.3, *ε* = 2.0×10^-3^%, *M* = 0.36, *SD* = 0.9 and *M* = −0.21, *SD* = 1.1, respectively). Furthermore, in order to determine if there was an impact of Val66Met genotype on the difference between familiarity and recollection across the LPC time window, we calculated the difference scores between items correctly identified as old with correct source-recognition (Hit-Hit), and items correctly identified as old with incorrect source-recognition (Hit-Miss). Using a Bayesian independent-samples *t*-test on these differences scores, we find moderate evidence for the null hypothesis that there is no difference in ERP amplitudes for items correctly identified as old with correct and incorrect source-recognition, across genotype groups (BF_01_= 3.7, *ε* = 2.0×10^-3^%, Val/Val: *M* = 0.16, *SD* = 0.77 and Met: *M* = 0.08, *SD* = 0.87, respectively).

Distributions for the scores of each group for each measure are shown in Figure 7. Posterior and prior distributions, as well as sequential analyses and robustness checks for these tests, are available in the Supplementary Material (Figures SM4:5).

**Figure 6.**
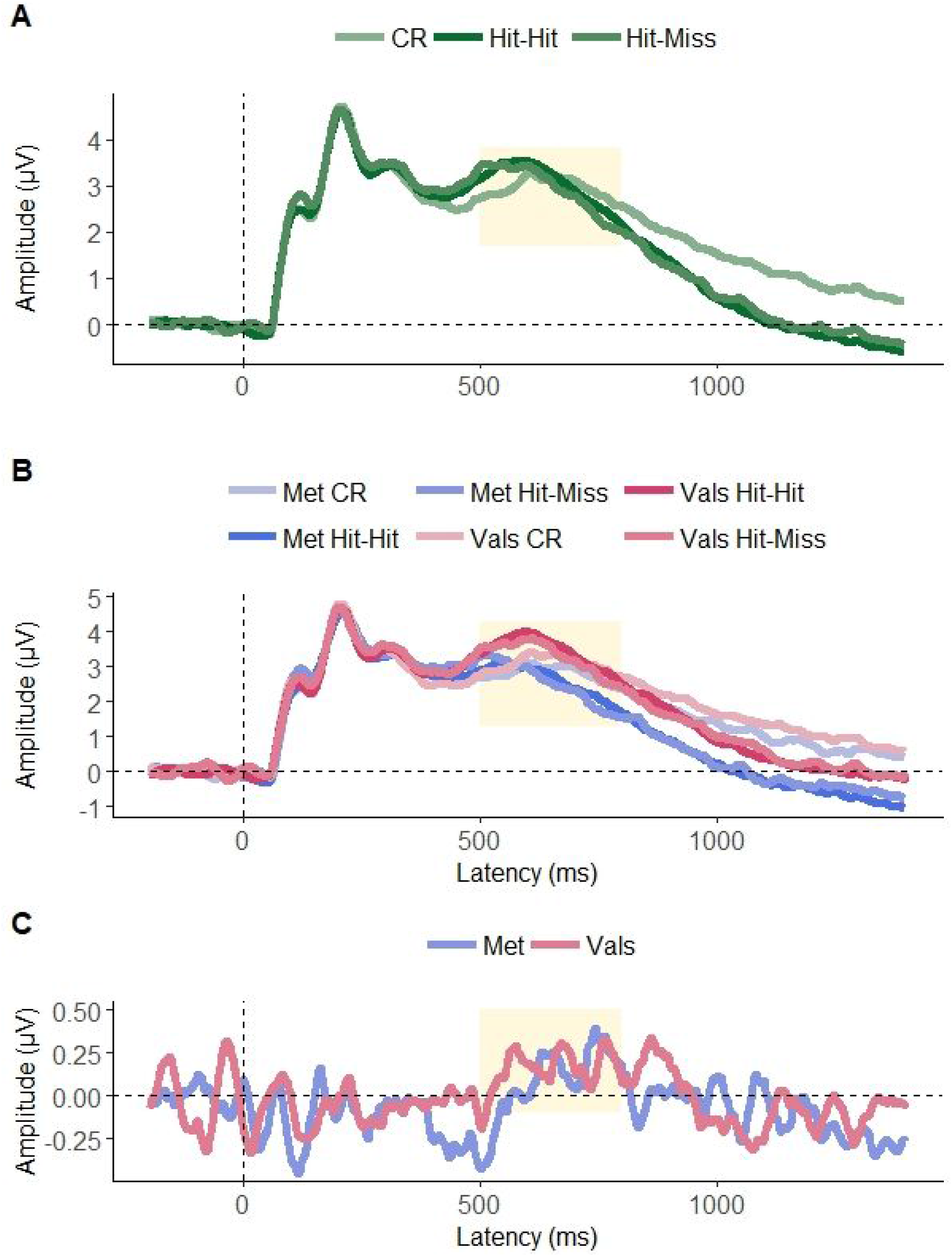
ERPs illustrating the electrophysiological responses to each of the correctly identified Old with correct source memory (Hit-Hits), correctly recognised old with incorrect source memory (Hit-Misses), and New (CR) items at the LPC electrode cluster. **A:** ERPs for the overall group showing no replication of previously reported Hit-Hit/Hit-Miss effects at posterior electrode sites. **B:** ERPs split by genotype demonstrating how each of Val/Val and Met allele individuals respond to different trial types. **C:** Difference waves calculated between ERPs elicited to Hit-Hit and ERPs elicited to Hit-Miss items. These are recorded at a cluster of electrodes centred around electrode 60, a left-ward, posterior, electrode. Time window of interest for the LPC component is highlighted in yellow (500-800ms).

**Figure 7.**
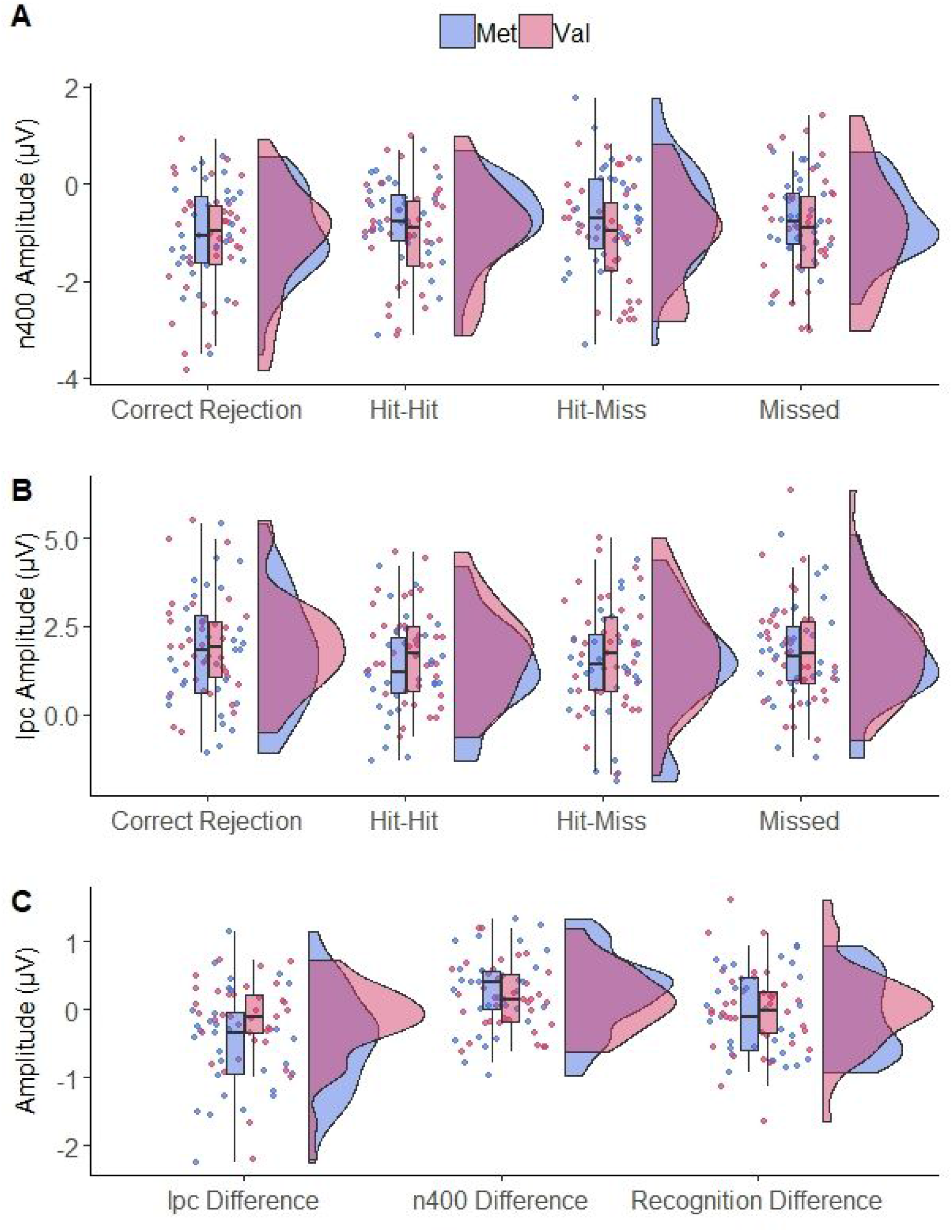
Distributions of the mean amplitude across the time windows of interest. **A:** Amplitude distributions for all four trial types, measured across the N400 time window (300-500ms post-stimulus). **B:** Amplitude distributions for all four trial types, measured across the LPC time window (500-800ms post-stimulus). **C:** Distributions of difference waves that were calculated for the three Bayesian independent-samples *t*-tests analysed in this chapter. These difference waves have been averaged across the specified time windows. In all instances Met allele carriers are depicted in blue, while Val/Val individuals are depicted in red.

### Behavioural-ERP correlations

We ran a Bayesian correlation analysis to relate all behavioural scores with average voltage within each condition for each time window of interest using the default stretched beta prior width of 1 (Morey & Rouder, 2018).

#### Correlations with the N400 time window

Mean voltage across the N400 time window for items correctly recognised as old (Hits), items correctly identified as new (Correct Rejections), and items that were incorrectly identified as new (Missed), were correlated with each of the unique behavioural measures (Hits, Correct Rejections, Item Accuracy, Source Accuracy, discriminability, response bias). We found anecdotal-moderate evidence for the null hypothesis that the ERP amplitudes in response to items correctly recognised as old were not correlated with our accuracy measures (Hits: *r* = −0.19, BF_01_ = 2.0, 95%CI = 0.05;-0.40; Correct Rejections: *r* = 0.20, BF_01_ = 1.8, 95%CI = 0.41;-0.04; Item Accuracy: *r* = −0.16, BF_01_ = 2.9, 95%CI = 0.08;-0.38; Source Accuracy: *r* = 0.06, BF_01_ = 6.0, 95%CI = 0.29;-0.18), or our discriminability measure (*r* = −0.09, BF_01_ = 5.2, 95%CI = 0.15;-0.31), and inconclusive evidence for whether response bias was correlated with average voltage (*r* = −0.25, BF_10_ = 1.2, 95%CI = −0.01;-0.46). Similar results were found for the average amplitude in response to new items correctly identified as new (Hits: *r* = −0.10, BF_01_ = 1.8, 95%CI = 0.04;-0.41; Correct Rejections: *r* = 0.20, BF_01_ = 1.8, 95%CI = 0.41;-0.04; Item Accuracy: *r* = −0.17, BF_01_ = 2.6, 95%CI = 0.07;-0.38; Source Accuracy: *r* = −0.00, BF_01_ = 6.6, 95%CI = 0.23;-0.24; discriminability: *r* = −0.10, BF_01_ = 4.9, 95%CI = 0.14;-0.32; response bias: *r* = −0.24, BF_10_ = 1.1, 95%CI = −0.00;-0.44), and old items incorrectly identified as new (Hits: *r* = −0.17, BF_01_ = 2.7, 95%CI = 0.07;-0.38; Correct Rejections: *r* = 0.06, BF_01_ = 6.0, 95%CI = 0.28;-0.18; Item Accuracy: *r* = −0.16, BF_01_= 2.9, 95%CI = 0.08;-0.38; Source Accuracy: *r* = −0.06, BF_01_ = 6.0, 95%CI = 0.28;-0.18; discriminability: *r* = −0.13, BF_01_= 3.7, 95%CI = 0.11;-0.35; response bias: *r* = −0.11, BF_01_ = 4.6, 95%CI = 0.13;-0.33).

#### Correlations with the LPC time window

Mean voltage across the LPC time window for items correctly recognised as old with incorrect source judgment (Hit-Miss), items correctly recognised as old with correct source judgment (Hit-Hit), and items correctly identified as new (Correct Rejections), were correlated with each of the unique behavioural measures ((Hits, Correct Rejections, Item Accuracy, Source Accuracy, discriminability, response bias). We found anecdotal-moderate evidence for the null hypothesis that the ERP amplitudes in response to items correctly recognised as old with incorrect source judgment were not correlated with our behavioural measures (Hits: *r* = −0.07, BF_01_ = 5.8, 95%CI = 0.17;-0.29; Correct Rejections: *r* = −0.15, BF_01_ = 3.2, 95%CI = 0.09;-0.37; Item Accuracy: *r* = −0.09, BF_01_ = 5.0, 95%CI = 0.15;-0.32; Source Accuracy: *r* = −0.02, BF_01_ = 6.5, 95%CI = 0.22;-0.25; discriminability: *r* = −0.14, BF_01_ = 3.6, 95%CI = 0.10;-0.36; response bias: *r* = 0.11, BF_01_ = 4.4, 95%CI = 0.33;-0.13). Consistent with these results, we also found moderate support for no correlation between items correctly identified as old with correct source judgment (Hits: *r* = −0.03, BF_01_ = 6.4, 95%CI = 0.20;-0.26; Correct Rejections: *r* = −0.17, BF_01_ = 2.6, 95%CI = 0.07;-0.39; Item Accuracy: *r* = −0.06, BF_01_ = 5.8, 95%CI = 0.18;-0.29; Source Accuracy: *r* = −0.06, BF_01_ = 6.0, 95%CI = 0.18;-0.28; discriminability: *r* = −0.12, BF_01_ = 4.3, 95%CI = 0.12;-0.34; response bias: *r* = 0.16, BF_01_ = 2.9, 95%CI = 0.38;-0.08).

## Discussion

In this study, we investigated the impact of the Val66Met SNP on the electrophysiological correlates of recognition memory. First, we aimed to replicate previous studies that reported a reduction in memory accuracy for carriers of the Met allele, when compared to Val/Val individuals (Egan et al., 2003; Hariri et al., 2003). We extended these reports by examining the relationship between Val66Met genotype and the two subprocesses of recognition memory; familiarity and recollection. In addition to these behavioural aims, we were also interested in examining whether the ERP components associated with each of these two recognition subprocesses are impacted by the Val66Met SNP, differentially. Previous research has indicated that familiarity processing is associated with a frontal negativity, the N400, while recollection is associated with a posterior positivity, the LPC. We, therefore, were interested in comparing the amplitudes of these ERPs in order to investigate how Val66Met genotype influences each of familiarity and recollection, electrophysiologically. Using a source memory task we compared the amplitudes of the ERPs of each condition (Hits, Hit-Hit, Correct Rejections) for each genotype group, across the time windows associated with the N400 and LPC. We hypothesised that given the reduction in memory performance reported in previous work, Met allele carriers would also have reduced ERP amplitudes on trials associated with accurate recognition. We further hypothesised that any differences in ERP amplitudes would be greatest for the LPC time window compared to the N400 time window, given that this is associated with recollection based processing.

Behaviourally, we replicated previous work reporting that Met allele carriers have reduced accuracy compared to Val/Val individuals on recognition memory tasks (Egan et al., 2003; Hariri et al., 2003; Spriggs et al., 2019). More specifically, we found anecdotal evidence in support of Val/Val individuals outperforming Met allele carriers on the item-recognition portion of our task. Our results provide some evidence for Val/Val individuals being better at recognising previously seen items, as well as identifying new items compared to carriers of the Met allele. This result is supported by other work also reporting greater recognition of previously seen items in Val homozygotes (Egan et al., 2003; Hariri et al., 2003). Interestingly, this pattern does not persist into the source-recognition subtask of our paradigm. Specifically, our results show evidence for the null hypothesis that Val/Val and Met allele carriers are equally accurate at remembering the encoding context of items that have been previously recognised. In line with this, our measure of discriminability (*Pr*) also provided evidence for Val/Val individuals being better able to discriminate old from new items. Finally, we also measured response bias (*Br*), using this measure we find support for the null, meaning that there does not appear to be any systematic difference in the response strategy of these groups. Together these findings suggest that the Met allele carriers of our sample are less accurate in recognising previously seen objects, however once recognised, they are equally able to recognise the encoding context of that previously seen object. We believe this is the first attempt to date to tease out the contributions that the Val66Met SNP makes to each of these subtasks of recognition. Given that the item-recognition subtask is a proxy measure of familiarity, and the source-recognition subtask is a proxy measure of recollection, our results seem to suggest that the Val66Met SNP influences the familiarity aspects of this source memory task only.

Electrophysiologically, we have partially replicated previous reports of a difference between ERP amplitudes for items that have been previously presented, compared to new items. These previous studies have reported that items which have been presented before tend to elicit a more positive ERP compared to items that are new (Rugg & Curran, 2007). We replicate this old/new effect for the N400, but not the LPC. More specifically, our results provide strong evidence for a difference in the ERP amplitudes to old versus new items presented during the item-recognition subtask. Interestingly, our results provide moderate support for the null hypothesis for the LPC time window, that there is no difference in the amplitude of ERPs elicited in response to old and new items. Additionally, recent work has reported that a dissociation in the LPC can be found between items that have been recognised with and without accurate source-recognition (Addante et al., 2012). For example, a dissociation in the LPC can be made between items that have been correctly recognised with correct source-recognition (Hit-Hit) and those that are correctly recognised with incorrect source-recognition (Hit-Miss). In an attempt to replicate this, we split our ERPs corresponding to correctly recognised objects into those with correct and incorrect source-recognition. However, our results provide moderate evidence for the null hypothesis of no difference between Hit-Hit and Hit-Miss ERP amplitudes across this time window, and therefore we did not replicate this dissociation.

Our primary interest was to assess the influence that the Val66Met genotype has on recognition related ERPs. In order to pursue this, we focused our analyses on the magnitude of the old/new effect. We calculated difference waves subtracting average amplitude in response to correctly identified new items from average amplitude in response to correctly recognised old items, for both the N400 and LPC components. As described above, the N400 is linked to item-recognition processing, while the LPC is linked to source-recognition processing. Our results show support for the null hypothesis that there was no difference in the size of the old/new effect for the N400 in the item-recognition subtask. However, we do find support for a difference in the size of the old/new effect for the LPC for the source-recognition subtask. That is, we found evidence for Val/Val individuals having larger differences in the amplitudes of ERPs produced in response to correctly recognised old items compared to correctly recognised new items. A final comparison was made between ERPs in response to correctly recognised old items with correct source-recognition and correctly recognised old items with incorrect source-recognition as a way to examine the impact of recollection over familiarity. In order to assess this, we subtracted the two associated ERPs. Our results show evidence for the null hypothesis that there was no difference in the size of this differenced ERP between the two genotypes. This is in direct contrast to our predictions as we originally hypothesised that differences in ERPs would be greatest for the LPC, given its link to recollection.

Interestingly, few EEG studies investigated the impact of the BDNF Val66Met SNP on memory. Most previous work comparing ERPs for these groups has focused on working memory (Schofield et al., 2009), attention (Getzmann et al., 2013), and error processing (Beste et al., 2010). While our results are not a direct comparison, we do notice one recurring relevant theme within these studies; all report an increase in ERP amplitude across their time windows of interest for Val/Val individuals compared to Met allele carriers. These increases in amplitude have been tied to various cognitive processes that are necessary for the respective tasks. Importantly, the increase in amplitude is not global across all regions of the ERPs of these studies, evidence that these modulations might be directly linked to differential task processing for individuals of these genotypes. Our results also reflect this pattern. Therefore, we propose this is evidence that our results are not merely the product of a general, global, processing difference between the two genotype groups. Consistent with these electrophysiological findings, several other studies interested in the impact of Val66Met genotype have focused on EEG oscillations and spectral power, rather than ERPs (Bachmann et al., 2012; Gatt et al., 2008; Geist, Dulka, Barnes, Totty, & Datta, 2017; Guindalini et al., 2014; Jones et al., 2017; Soltész et al., 2014). Generally, these studies have also found dampened electrophysiological activity in Met allele carriers compared to Val/Val individuals.

Our results add to a large body of literature that describes the functional impact of the BDNF Val66Met SNP (Egan et al., 2003; Hariri et al., 2003; Ho et al., 2006). Given that BDNF is known to be secreted in response to cellular activity, and that the Met allele is furthermore associated with reduced secretion (Chen et al., 2004) and impaired movement of the protein within the cell (Farhadi et al., 2000), Met allele carriers have lower concentrations of available BDNF protein. There are clear demonstrations of the importance of the BDNF protein to long-term potentiation (Poo, 2001), a cellular mechanism underpinning memory. Therefore, we speculate that this reduction in available BDNF could underpin the memory and ERP differences we detected. This would mean our results could be an illustration of the immediate functional impact of the BDNF SNP rather than evidence of any long-term, developmentally derived, influence.

More specifically, we observed a behavioural disadvantage of the Met allele during the item-recognition—but not the source-recognition—subtask of our overall source memory task. This observation is consistent across judgments and item types (i.e., correctly recognising old items, correctly identifying new items, and overall item-recognition accuracy), likely a result of an increased discriminability, which we also found for our Val/Val individuals. Importantly, these patterns were not the result of a change in response strategy between the two groups.

Interestingly, our ERP observations show a difference between the Val66Met genotype groups that is specific to the source-recognition subtask and not the item-recognition subtask. While at first these results seem to be contradictory, it is possible they are each reflecting an overall difference in the old/new effect between our two groups, and are therefore complementary findings. Evidence for this comes from the fact that we did not replicate the late dissociation between familiarity and recollection across the LPC component, as observed in previous research (Addante et al., 2012). Despite this, our results are supported by several other groups that have also failed to dissociate familiarity and recollection during this time window (Gold et al., 2006; Wais, Wixted, Hopkins, & Squire, 2006). Overall, we report a behavioural impact of the Met allele for the item-recognition subtask, while we are only able to observe the old/new difference, electrophysiologically, at the LPC time point.

One possible alternative explanation for our results is that structural changes between the genotype groups could result in electrophysiological differences. For example, it has been previously reported that there are differences in cortical thickness (Nemoto et al., 2006; Yang et al., 2012), as well as subcortical volumetric differences (Bueller et al., 2006; Pezawas et al., 2004; Szeszko et al., 2005) between these groups. Given that the EEG signal is known to be highly impacted by underlying cortical structure, this could be an issue if there are systematic differences in brain morphometry between Val/Val individuals and Met carriers. While the current study does not directly investigate this question, a subset of these individuals also participated in an MRI study that aimed to investigate possible structural differences associated with the Val66Met SNP (McKay et al., 2019). This study found no detectable differences in ROI or global analyses for either grey- or white-matter measures, and therefore, instils some confidence that our results might not be an artefact of large structural variations in these groups. Furthermore, the observation that our results are constrained to the LPC time window also suggests that we are measuring something specific, rather than a diffuse electrophysiological difference.

There are some significant limitations to consider when evaluating our results. First, our results showed no evidence for a correlation between the size of the old/new effect at either component (N400 or LPC), and the accuracy on either the item-recognition or source-recognition subtasks. This is problematic and limits our ability to draw strong conclusions about what this increase in amplitude across the recollection-relevant time window might reflect. Furthermore, we find behavioural differences between the two genotype groups that are directly linked to the item-recognition subtask, while our ERP differences occur at the later LPC, associated with the source-recognition subtask. While this might seem incongruous, it is possible that both the behavioural and ERP results are reflecting an overall influence of the Val66Met SNP on the old/new effect (i.e., general recognition), rather than any impact specific to either familiarity or recollection. A further related limitation is that we were unable to replicate the findings of Addante et al. (2012) who found dissociations between familiarity and recollection-related ERPs at the late time window. One reason for our divergent results could be that the original paper used confidence ratings to ensure they were able to contrast only items that were remembered with the highest confidence, as their recollection sample. Due to time constraints, we were unable to probe our participants for confidence ratings during our recognition trials, therefore, our study was not an exact replication of the original experiment. This points to the importance of contrasting familiarity with only high confidence recollection, an observation highlighted in several key papers (Mitchell & Johnson, 2009; Woodruff et al., 2006; Yu & Rugg, 2010).

A final major limitation we wish to acknowledge relates to the source-recognition memory task itself. This task involves participants making two binary decisions, one regarding item-recognition (“old” or “new”) and one regarding source-recognition (“manmade” or “box”). These decisions do not necessarily reflect the exact nature of the familiarity and recollection processes that we associate them with. While one option could be to add confidence ratings and make each decision a 5-point scale, it is also possible that another task would be better suited to dissociate familiarity and recollection. For example, the remember-know task has also previously been used to dissociate familiarity from recollection (Duarte, Ranganath, Winward, Hayward, & Knight, 2004; Mollison & Curran, 2012; Vilberg, Moosavi, & Rugg, 2006). While most previous studies have found that the source-recognition subtask is generally associated with recollection, there are several examples where researchers have found familiarity to be equally associated with source memory (Duarte, Ranganath, Trujillo, & Knight, 2006; Hicks, Marsh, & Ritschel, 2002). These observations are important as they illustrate that we may not be able to entirely dissociate familiarity from recollection, using the source memory task. However, one important reason why the source-recognition task was chosen for this study was its unique ability to potentially dissociate familiarity and recollection within single trials, thus facilitating its implementation in an EEG design.

Despite these limitations, our results do clearly extend previous studies by providing behavioural evidence that differences between individuals with and without the Met allele, might be related to the familiarity subcomponent of recognition. This finding could be underpinned by differences in discriminability, which might reflect that Val/Val individuals are better able to discriminate between old and new items. Supporting this behavioural finding, we also detect electrophysiological differences between these groups, however, this seems to be constrained to the late recognition component, commonly linked to the recollection subcomponent of recognition. Given that other research groups have argued familiarity to also modulate the LPC, we propose our findings might best be described as reflecting the impact of the Val66Met SNP on general recognition, or potentially on familiarity-related recognition. Importantly, these behavioural and electrophysiological results are reported in the absence of any major structural changes within the brain (see McKay et al., 2019). We, therefore, propose that these results are evidence of a functional influence of the Val66Met SNP on recognition memory.

## Data availability statement

Due to restrictions imposed by the University of Auckland Human Participants Ethics Committee, participant-level genetic and EEG data are not available for sharing. However, aggregate data and research code is available upon request, by contacting the corresponding author.

## References

Addante, R. J., Ranganath, C., & Yonelinas, A. P. (2012). Examining ERP correlates of recognition memory: evidence of accurate source recognition without recollection. NeuroImage, 62(1), 439–450. https://doi.org/10.1016/j.neuroimage.2012.04.031

Aggleton, J. P., & Brown, M. W. (1999). Episodic memory, amnesia, and the hippocampal–anterior thalamic axis. The Behavioral and Brain Sciences, 22(3), 425–444. Retrieved from https://www.cambridge.org/core/journals/behavioral-and-brain-sciences/article/episodic-memory-amnesia-and-the-hippocampalanterior-thalamic-axis/810189AC34B388075020FA6E6C61F000

Aggleton, J. P., Dumont, J. R., & Warburton, E. C. (2011). Unraveling the contributions of the diencephalon to recognition memory: a review. Learning & Memory, 18(6), 384–400. https://doi.org/10.1101/lm.1884611

Bachmann, V., Klein, C., Bodenmann, S., Schäfer, N., Berger, W., Brugger, P., & Landolt, H.-P. (2012). The BDNF Val66Met polymorphism modulates sleep intensity: EEG frequency- and state-specifìcity. Sleep, 35(3), 335–344. https://doi.org/10.5665/sleep.1690

Beste, C., Kolev, V., Yordanova, J., Domschke, K., Falkenstein, M., Baune, B. T., & Konrad, C. (2010). The role of the BDNF Val66Met polymorphism for the synchronization of error-specific neural networks. The Journal of Neuroscience: The Official Journal of the Society for Neuroscience, 30(32), 10727–10733. https://doi.org/10.1523/JNEUROSCI.2493-10.2010

Brodeur, M. B., Dionne-Dostie, E., Montreuil, T., & Lepage, M. (2010). The Bank of Standardized Stimuli (BOSS), a new set of 480 normative photos of objects to be used as visual stimuli in cognitive research. PloS One, 5(5), el0773. https://doi.org/10.1371/journal.pone.0010773

Brodeur, M. B., Guérard, K., & Bouras, M. (2014). Bank of Standardized Stimuli (BOSS) phase II: 930 new normative photos. PloS One, 9(9), el06953. https://doi.org/10.1371/journal.pone.0106953

Brown, M. W., & Xiang, J. Z. (1998). Recognition memory: neuronal substrates of the judgement of prior occurrence. Progress in Neurobiology, 55(2), 149–189. https://doi.org/10.1016/S0301-0082(98)00002-1

Bueller, J. A., Aftab, M., Sen, S., Gomez-Hassan, D., Burmeister, M., & Zubieta, J.-K. (2006). BDNF Val66Met allele is associated with reduced hippocampal volume in healthy subjects. Biological Psychiatry, 59(9), 812–815. https://doi.org/10.1016/j.biopsych.2005.09.022

Cansino, S., & Trejo-Morales, P. (2008). Neurophysiology of successful encoding and retrieval of source memory. Cognitive, Affective & Behavioral Neuroscience, 5(1), 85–98. https://doi.org/10.3758/CABN.8.1.85

Chen, Z.-Y., Patel, P. D., Sant, G., Meng, C.-X., Teng, K. K., Hempstead, B. L., & Lee, F. S. (2004). Variant brain-derived neurotrophic factor (BDNF) (Met66) alters the intracellular trafficking and activity-dependent secretion of wild-type BDNF in neurosecretory cells and cortical neurons. The Journal of Neuroscience: The Official Journal of the Society for Neuroscience, 24(18), 4401–4411. https://doi.org/10.1523/JNEUROSCI.0348-04.2004

Curran, T. (2000). Brain potentials of recollection and familiarity. Memory & Cognition, 28(6), 923–938. https://doi.org/10.3758/BF03209340

Delorme, A., & Makeig, S. (2004). EEGLAB: an open source toolbox for analysis of single-trial EEG dynamics including independent component analysis. Journal of Neuroscience Methods, 134(1), 9–21. https://doi.org/10.1016/j.jneumeth.2003.10.009

Dempster, E., Toulopoulou, T., McDonald, C., Bramon, E., Walshe, M., Filbey, F., … Collier, D. A. (2005). Association between BDNF val66 met genotype and episodic memory. American Journal of Medical Genetics. Part B, Neuropsychiatric Genetics: The Official Publication of the International Society of Psychiatric Genetics, 134B(1), 73–75. https://doi.org/10.1002/ajmg.b.30150

Duarte, A., Ranganath, C., Trujillo, C., & Knight, R. T. (2006). Intact recollection memory in high-performing older adults: ERP and behavioral evidence. Journal of Cognitive Neuroscience, 18(1), 33–47. https://doi.org/10.1162/089892906775249988

Duarte, A., Ranganath, C., Winward, L., Hayward, D., & Knight, R. T. (2004). Dissociable neural correlates for familiarity and recollection during the encoding and retrieval of pictures. Brain Research. Cognitive Brain Research, 18(3), 255–272. Retrieved from https://www.ncbi.nlm.nih.gov/pubmed/14741312

Düzel, E., Yonelinas, A. P., Mangun, G. R., Heinze, H.-J., & Tulving, E. (1997). Event-related brain potential correlates of two states of conscious awareness in memory. Proceedings of the National Academy of Sciences of the United States of America, 94(11), 5973–5978. https://doi.org/10.1073/pnas.94.11.5973

Egan, M. F., Kojima, M., Callicott, J. H., Goldberg, T. E., Kolachana, B. S., Bertolino, A., … Weinberger, D. R. (2003). The BDNF val66met polymorphism affects activity-dependent secretion of BDNF and human memory and hippocampal function. Cell, 112(2), 257–269. https://doi.org/10.1016/S0092-8674(03)00035-7

Eichenbaum, H., Yonelinas, A. P., & Ranganath, C. (2007). The medial temporal lobe and recognition memory. Annual Review of Neuroscience, 30, 123–152. https://doi.org/10.l146/annurev.neuro.30.051606.094328

Farhadi, H. F., Mowla, S. J., Petrecca, K., Morris, S. J., Seidah, N. G., & Murphy, R. A. (2000). Neurotrophin-3 sorts to the constitutive secretory pathway of hippocampal neurons and is diverted to the regulated secretory pathway by coexpression with brain-derived neurotrophic factor. The Journal of Neuroscience: The Official Journal of the Society for Neuroscience, 20(11), 4059–4068. Retrieved from https://www.ncbi.nlm.nih.gov/pubmed/10818141

Friedman, D., & Johnson, R., Jr. (2000). Event-related potential (ERP) studies of memory encoding and retrieval: a selective review. Microscopy Research and Technique, 51(1), 6–28. https://doi.org/3.0.CO;2-R”>10.1002/1097-0029(20001001)51:1<6::AID-JEMT2>3.0.CO;2-R

Gajewski, P. D., Hengstler, J. G., Golka, K., Falkenstein, M., & Beste, C. (2012). The Met-genotype of the BDNF Val66Met polymorphism is associated with reduced Stroop interference in elderly. Neuropsychologia, 50(14), 3554–3563. https://doi.org/10.1016/j.neuropsychologia.2012.09.042

Gatt, J. M., Kuan, S. A., Dobson-Stone, C., Paul, R. H., Joffe, R. T., Kemp, A. H., … Williams, L. M. (2008). Association between BDNF Val66Met polymorphism and trait depression is mediated via resting EEG alpha band activity. Biological Psychology, 79(2), 275–284. https://doi.org/10.1016/j.biopsycho.2008.07.004

Geist, P. A., Dulka, B. N., Barnes, A., Totty, M., & Datta, S. (2017). BNDF heterozygosity is associated with memory deficits and alterations in cortical and hippocampal EEG power. Behavioural Brain Research, 332, 154–163. https://doi.org/10.1016/j.bbr.2017.05.039

Getzmann, S., Gajewski, P. D., Hengstler, J. G., Falkenstein, M., & Beste, C. (2013). BDNF Val66Met polymorphism and goal-directed behavior in healthy elderly - evidence from auditory distraction. Neuroimage, 64, 290–298. https://doi.org/10.1016/j.neuroimage.2012.08.079

Gold, J. J., Smith, C. N., Bayley, P. J., Shrager, Y., Brewer, J. B., Stark, C. E. L., … Squire, L. R. (2006). Item memory, source memory, and the medial temporal lobe: concordant findings from fMRI and memory-impaired patients. Proceedings of the National Academy of Sciences of the United States of America, 103(24), 9351–9356. https://doi.org/10.1073/pnas.0602716103

Gruber, M. J., & Otten, L. J. (2010). Voluntary control over prestimulus activity related to encoding. The Journal of Neuroscience: The Official Journal of the Society for Neuroscience, 30(29), 9793–9800. https://doi.org/10.1523/JNEUROSCI.0915-10.2010

Guindalini, C., Mazzotti, D. R., Castro, L. S., D’Aurea, C. V. R., Andersen, M. L., Poyares, D., … Tufik, S. (2014). Brain-derived neurotrophic factor gene polymorphism predicts interindividual variation in the sleep electroencephalogram. Journal of Neuroscience Research, 92(8), 1018–1023. https://doi.org/10.1002/jnr.23380

Hariri, A. R., Goldberg, T. E., Mattay, V. S., Kolachana, B. S., Callicott, J. H., Egan, M. F., & Weinberger, D. R. (2003). Brain-derived neurotrophic factor val66met polymorphism affects human memory-related hippocampal activity and predicts memory performance. The Journal of Neuroscience: The Official Journal of the Society for Neuroscience, 23(11), 6690–6694. https://doi.org/10.1523/JNEUROSCI.23-17-06690.2003

Henson, R. N., Rugg, M. D., Shallice, T., Josephs, O., & Dolan, R. J. (1999). Recollection and familiarity in recognition memory: an event-related functional magnetic resonance imaging study. The Journal of Neuroscience: The Official Journal of the Society for Neuroscience, 79(10), 3962–3972. https://doi.org/10.1523/JNEUROSCI.19-10-03962.1999

Hicks, J. L., Marsh, R. L., & Ritschel, L. (2002). The role of recollection and partial information in source monitoring. Journal of Experimental Psychology. Learning, Memory, and Cognition, 28(3), 503–508. Retrieved from https://www.ncbi.nlm.nih.gov/pubmed/12018502

Hintzman, D. L., Caulton, D. A., & Levitin, D. J. (1998). Retrieval dynamics in recognition and list discrimination: further evidence of separate processes of familiarity and recall. Memory & Cognition, 26(3), 449–462. Retrieved from https://www.ncbi.nlm.nih.gov/pubmed/9610117

Hiramoto, R., Kanayama, N., Nakao, T., Matsumoto, T., Konishi, H., Sakurai, S., … Yamawaki, S. (2017). BDNF as a possible modulator of EEG oscillatory response at the parietal cortex during visuo-tactile integration processes using a rubber hand. Neuroscience Research, 124, 16–24. https://doi.org/10.1016/j.neures.2017.05.006

Ho, B.-C., Milev, P., O’Leary, D. S., Librant, A., Andreasen, N. C., & Wassink, T. H. (2006). Cognitive and magnetic resonance imaging brain morphometric correlates of brain-derived neurotrophic factor Val66Met gene polymorphism in patients with schizophrenia and healthy volunteers. Archives of General Psychiatry, 63(1), 731–740. https://doi.org/10.1001/archpsyc.63.7.731

Hoppstädter, M., Baeuchl, C., Diener, C., Flor, H., & Meyer, P. (2015). Simultaneous EEG-fMRI reveals brain networks underlying recognition memory ERP old/new effects. NeuroImage, 116, 112–122. https://doi.org/10.1016/j.neuroimage.2015.05.026

Johnson, M. K., Hashtroudi, S., & Lindsay, D. S. (1993). Source monitoring. Psychological Bulletin, 114(1), 3–28. https://doi.org/10.1037/0033-2909.114.1.3

Jones, N. C., Hudson, M., Foreman, J., Rind, G., Hill, R., Manning, E. E., & van den Buuse, M. (2017). Brain-derived neurotrophic factor haploinsuffìciency impairs high-frequency cortical oscillations in mice. The European Journal of Neuroscience. https://doi.org/10.1111/ejn.13722

Kennedy, K. M., Reese, E. D., Horn, M. M., Sizemore, A. N., Unni, A. K., Meerbrey, M. E., … Rodrigue, K. M. (2015). BDNF val66met polymorphism affects aging of multiple types of memory. Brain Research, 1612, 104–117. https://doi.org/10.1016/j.brainres.2014.09.044

Komulainen, P., Pedersen, M., Hänninen, T., Bruunsgaard, H., Lakka, T. A., Kivipelto, M., … Rauramaa, R. (2008). BDNF is a novel marker of cognitive function in ageing women: the DR’s EXTRA Study. Neurobiology of Learning and Memory, 90(4), 596–603. https://doi.org/10.1016/j.nlm.2008.07.014

Lamb, Y. N., Thompson, C. S., McKay, N. S., Waldie, K. E., & Kirk, I. J. (2015). The brain-derived neurotrophic factor (BDNF) val66met polymorphism differentially affects performance on subscales of the Wechsler Memory Scale - Third Edition (WMS-III). Frontiers in Psychology, 6, 1212. https://doi.org/10.3389/fpsyg.2015.01212

Leynes, P. A., Landau, J., Walker, J., & Addante, R. J. (2005). Event-related potential evidence for multiple causes of the revelation effect. Consciousness and Cognition, 14(2), 327–350. https://doi.org/10.1016/j.concog.2004.08.005

Luck, S. J. (2014). An introduction to the event-related potential technique. MIT press.

McKay, N. S., Moreau, D., Henare, D. T., & Kirk, I. J. (2019). The brain-derived neurotrophic factor Val66Met genotype does not influence the grey or white matter structures underlying recognition memory. NeuroImage. https://doi.org/10.1016/j.neuroimage.2019.03.072

Mitchell, K. J., & Johnson, M. K. (2009). Source Monitoring 15 Years Later: What Have We Learned From fMRI About the Neural Mechanisms of Source Memory? Psychological Bulletin, 735(4), 638–677. https://doi.org/10.1037/a0015849

Mollison, M. V., & Curran, T. (2012). Familiarity in source memory. Neuropsychologia, 50(11), 2546–2565. https://doi.org/10.1016/j.neuropsychologia.2012.06.027

Morey, R., & Rouder, J. (2018). BayesFactor: Computation of Bayes Factors for Common Designs. R package version 0.9.12-4.2. Retrieved from https://CRAN.R-project.org/package=BayesFactor

Mullen, T. (2012). CleanLine EEGLAB plugin. San Diego, CA: Neuroimaging Informatics Toolsand Resources Clearinghouse (NITRC).

Nemoto, K., Ohnishi, T., Mori, T., Moriguchi, Y., Hashimoto, R., Asada, T., & Kunugi, H. (2006). The Val66Met polymorphism of the brain-derived neurotrophic factor gene affects age-related brain morphology. Neuroscience Letters, 397(1-2), 25–29. https://doi.org/10.1016/j.neulet.2005.11.067

Payton, A. (2006). Investigating cognitive genetics and its implications for the treatment of cognitive deficit. Genes, Brain, and Behavior, 5 Suppl 1, 44–53. https://doi.org/10.1111/j.1601-183X.2006.00194.x

Pezawas, L., Verchinski, B. A., Mattay, V. S., Callicott, J. H., Kolachana, B. S., Straub, R. E., … Weinberger, D. R. (2004). The brain-derived neurotrophic factor val66met polymorphism and variation in human cortical morphology. The Journal of Neuroscience: The Official Journal of the Society for Neuroscience, 24(45), 10099–10102. https://doi.org/10.1523/JNEUROSCI.2680-04.2004

Poo, M. M. (2001). Neurotrophins as synaptic modulators. Nature Reviews. Neuroscience, 2(1), 24–32. https://doi.org/10.1038/35049004

R Core Team. (2019). R: A language and environment for statistical computing (Version 3.6.0). Retrieved from https://www.R-project.org/

Rugg, M. D., & Curran, T. (2007). Event-related potentials and recognition memory. Trends in Cognitive Sciences, 11(6), 251–257. https://doi.org/10.1016/j.tics.2007.04.004

Schofield, P. R., Williams, L. M., Paul, R. H., Gatt, J. M., Brown, K., Luty, A., … Gordon, E. (2009). Disturbances in selective information processing associated with the BDNF Val66Met polymorphism: evidence from cognition, the P300 and fronto-hippocampal systems. Biological Psychology, 80(2), 176–188. https://doi.org/10.1016/j.biopsycho.2008.09.001

Soltész, F., Suckling, J., Lawrence, P., Tait, R., Ooi, C., Bentley, G., … Nathan, P. J. (2014). Identification of BDNF sensitive electrophysiological markers of synaptic activity and their structural correlates in healthy subjects using a genetic approach utilizing the functional BDNF Val66Met polymorphism. PloS One, 9(4), e95558. https://doi.org/10.1371/journal.pone.0095558

Spriggs, M. J., Thompson, C. S., Moreau, D., McNair, N. A., Wu, C. C., Lamb, Y. N., … Kirk, I. J. (2019). Human Sensory LTP Predicts Memory Performance and Is Modulated by the BDNF Val66Met Polymorphism. Frontiers in Human Neuroscience, 13, 22. https://doi.org/10.3389/fnhum.2019.00022

Szeszko, P. R., Lipsky, R., Mentschel, C., Robinson, D., Gunduz-Bruce, H., Sevy, S., … Malhotra, A. K. (2005). Brain-derived neurotrophic factor val66met polymorphism and volume of the hippocampal formation. Molecular Psychiatry, 10(7), 631–636. https://doi.org/10.1038/sj.mp.4001656

Vilberg, K. L., Moosavi, R. F., & Rugg, M. D. (2006). The relationship between electrophysiological correlates of recollection and amount of information retrieved. Brain Research, 1122(1), 161–170. https://doi.org/10.1016/j.brainres.2006.09.023

Voss, J. L., & Paller, K. A. (2007). Neural correlates of conceptual implicit memory and their contamination of putative neural correlates of explicit memory. Learning & Memory, 14(4), 259–267. https://doi.org/10.1101/lm.529807

Wais, P. E., Wixted, J. T., Hopkins, R. O., & Squire, L. R. (2006). The hippocampus supports both the recollection and the familiarity components of recognition memory. Neuron, 49(3), 459–466. https://doi.org/10.1016/j.neuron.2005.12.020

Wilding, E. L., & Rugg, M. D. (1996). An event-related potential study of recognition memory with and without retrieval of source. Brain: A Journal of Neurology, 119 (Pt 3), 889–905. https://doi.org/10.1093/brain/119.3.889

Winkler, I., Debener, S., Müller, K. R., & Tangermann, M. (2015). On the influence of high-pass filtering on ICA-based artifact reduction in EEG-ERP. 2015 37th Annual International Conference of the IEEE Engineering in Medicine and Biology Society (EMBC), 4101–4105. https://doi.org/10.1109/EMBC.2015.7319296

Wixted, J. T. (2007). Dual-process theory and signal-detection theory of recognition memory. Psychological Review, 114(1), 152–176. https://doi.org/10.1037/0033-295X.114.1.152

Woodruff, C. C., Hayama, H. R., & Rugg, M. D. (2006). Electrophysiological dissociation of the neural correlates of recollection and familiarity. Brain Research, 1100(1), 125–135. https://doi.org/10.1016/j.brainres.2006.05.019

Yang, X., Liu, P., Sun, J., Wang, G., Zeng, F., Yuan, K., … Tian, J. (2012). Impact of brain-derived neurotrophic factor Val66Met polymorphism on cortical thickness and voxel-based morphometry in healthy Chinese young adults. PloS One, 7(6), e37777. https://doi.org/10.1371/journal.pone.0037777

Yonelinas, A. P. (1994). Receiver-operating characteristics in recognition memory: evidence for a dual-process model. Journal of Experimental Psychology. Learning, Memory, and Cognition, 20(6), 1341–1354. Retrieved from https://www.ncbi.nlm.nih.gov/pubmed/7983467

Yonelinas, A. P. (2002). The Nature of Recollection and Familiarity: A Review of 30 Years of Research. Journal of Memory and Language, 46(3), 441–517. https://doi.org/10.1006/jmla.2002.2864

Yonelinas, A. P., Kroll, N. E. A., Dobbins, I. G., Lazzara, M., & Knight, R. T. (1999). The neural substrates of recollection and familiarity. The Behavioral and Brain Sciences, 22(3), 468–469. https://doi.org/10.1017/S0140525X9946203X

Yonelinas, A. P., Otten, L. J., Shaw, K. N., & Rugg, M. D. (2005). Separating the brain regions involved in recollection and familiarity in recognition memory. The Journal of Neuroscience: The Official Journal of the Society for Neuroscience, 25(11), 3002–3008. https://doi.org/10.1523/JNEUROSCI.5295-04.2005

Yonelinas, A. P., & Parks, C. M. (2007). Receiver operating characteristics (ROCs) in recognition memory: a review. Psychological Bulletin, 133(5), 800–832. https://doi.org/10.1037/0033-2909.133.5.800

Yu, S. S., & Rugg. (2010). Dissociation of the electrophysiological correlates of familiarity strength and item repetition. Brain Research, 1320, 74–84. https://doi.org/10.1016/j.brainres.2009.12.071

