## Supplementary Material for "Investigating the Influence of the Brain-Derived Neurotrophic Factor Val66Met Single Nucleotide Polymorphism on Familiarity and Recollection Event-Related Potentials"

### Supporting Bayesian Plots

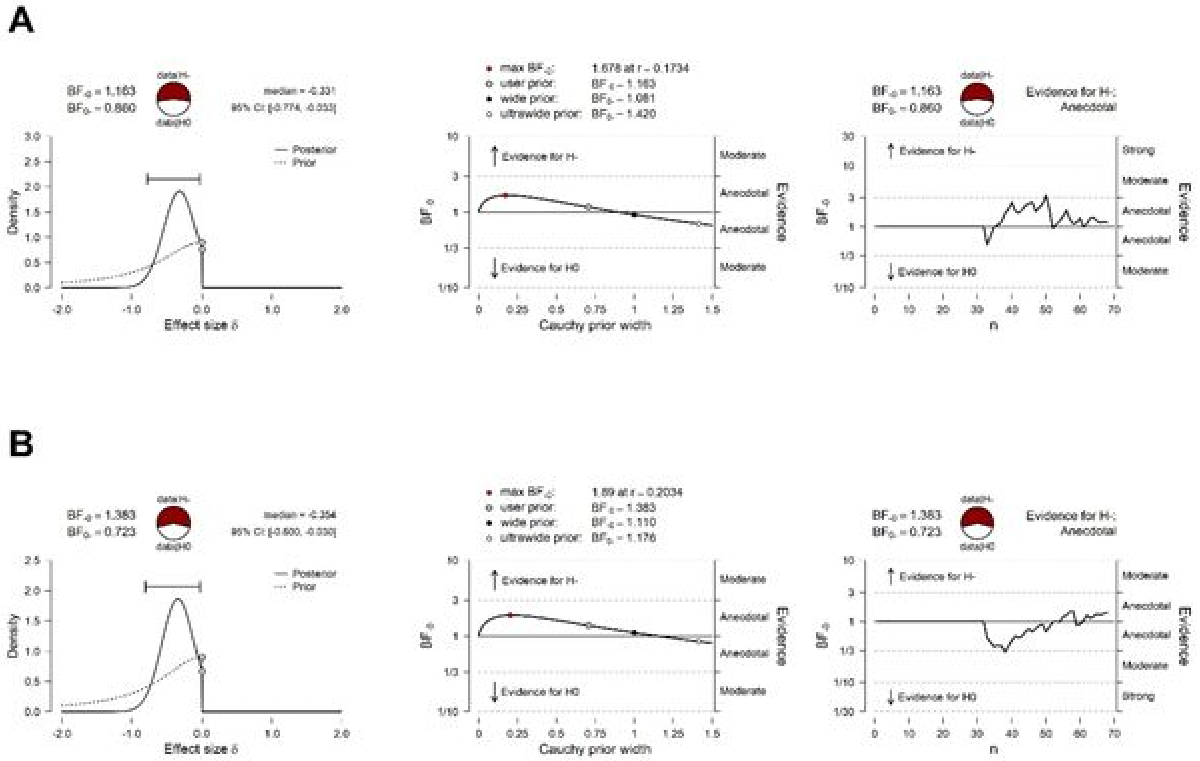


**Figure SM.1**. **Supporting plots for the hit and correct rejection measures**. Panel A: Prior and posterior distributions, robustness checks, and sequential analysis for the *t*-test conducted on the hit scores. Panel B: Prior and posterior distributions, robustness checks, and sequential analysis for the *t*-test conducted on the correct rejection scores. Robustness checks show how the resulting Bayes factor changes across a wide range of priors. Sequential analysis depict the evidence for each of the null and alternative hypotheses, as sample size increases. Accompanying Bayes factors and the credible interval are also displayed.

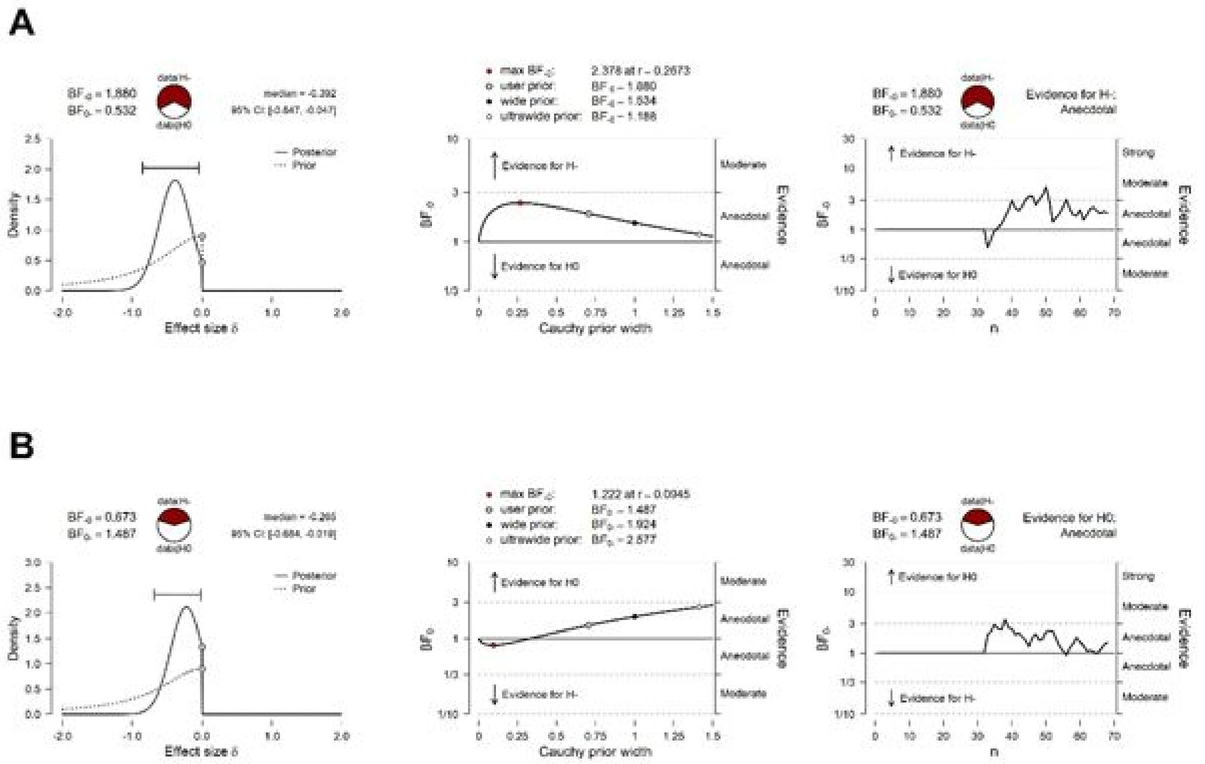


**Figure SM.2**. **Supporting plots for the item recognition and source recognition measures**. Panel A: Prior and posterior distributions, robustness checks, and sequential analysis for the *t*-test conducted on the item recognition scores. Panel B: Prior and posterior distributions, robustness checks, and sequential analysis for the *t*-test conducted on the source recognition scores. Robustness checks show how the resulting Bayes factor changes across a wide range of priors. Sequential analysis depict the evidence for each of the null and alternative hypotheses, as sample size increases. Accompanying Bayes factors and the credible interval are also displayed.

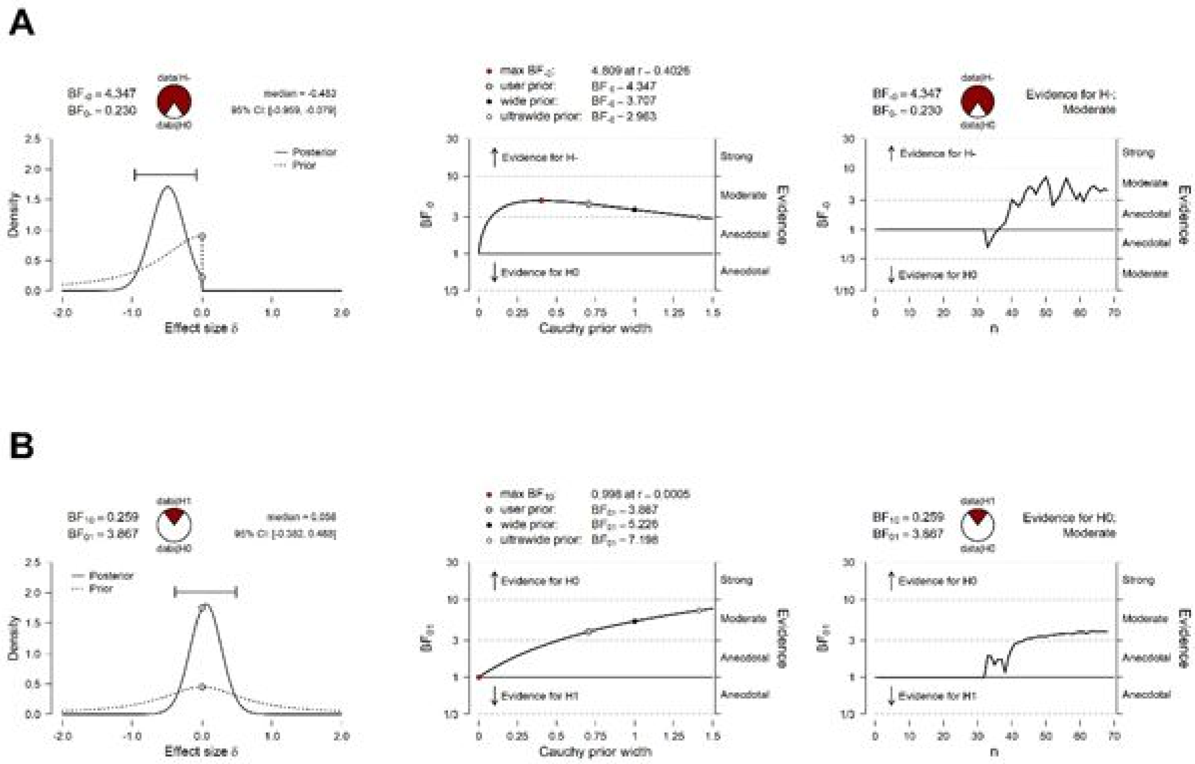


**Figure SM.3**. **Supporting plots for the discriminability and response bias measures**. Panel A: Prior and posterior distributions, robustness checks, and sequential analysis for the *t*-test conducted on the discriminability values. Panel B: Prior and posterior distributions, robustness checks, and sequential analysis for the *t*-test conducted on the response bias scores. Robustness checks show how the resulting Bayes factor changes across a wide range of priors. Sequential analysis depict the evidence for each of the null and alternative hypotheses, as sample size increases. Accompanying Bayes factors and the credible interval are also displayed.

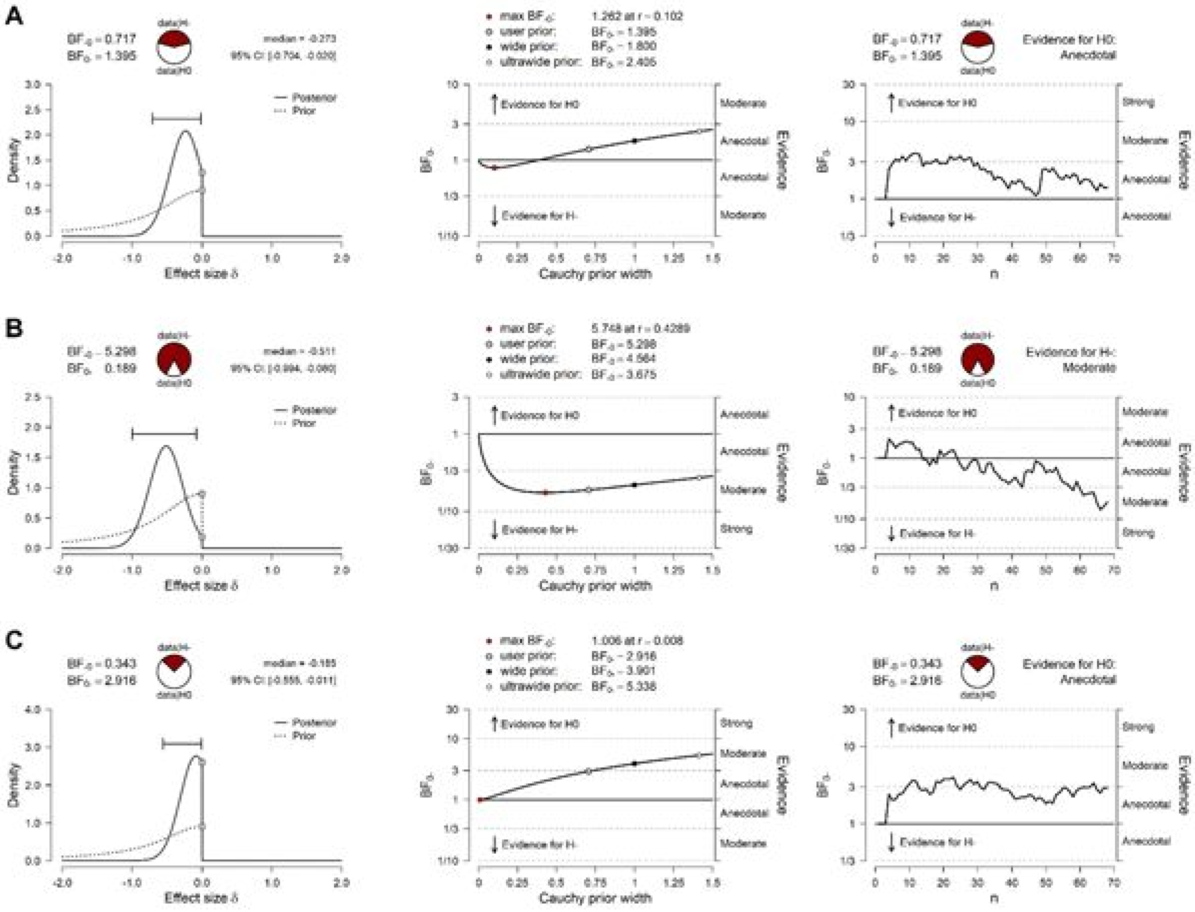


**Figure SM.4**. **Supporting plots for the EEG measures**. Panel A: Prior and posterior distributions, robustness checks, and sequential analysis for the *t*-test conducted on the N400 amplitude values. Panel B: Prior and posterior distributions, robustness checks, and sequential analysis for the *t*-test conducted on the LPC amplitude values. Panel C: Prior and posterior distributions, robustness checks, and sequential analysis for the *t*-test conducted on the hit-hit and hit-miss LPC amplitude values. Robustness checks show how the resulting Bayes factor changes across a wide range of priors. Sequential analysis depict the evidence for each of the null and alternative hypotheses, as sample size increases. Accompanying Bayes factors and the credible interval are also displayed.

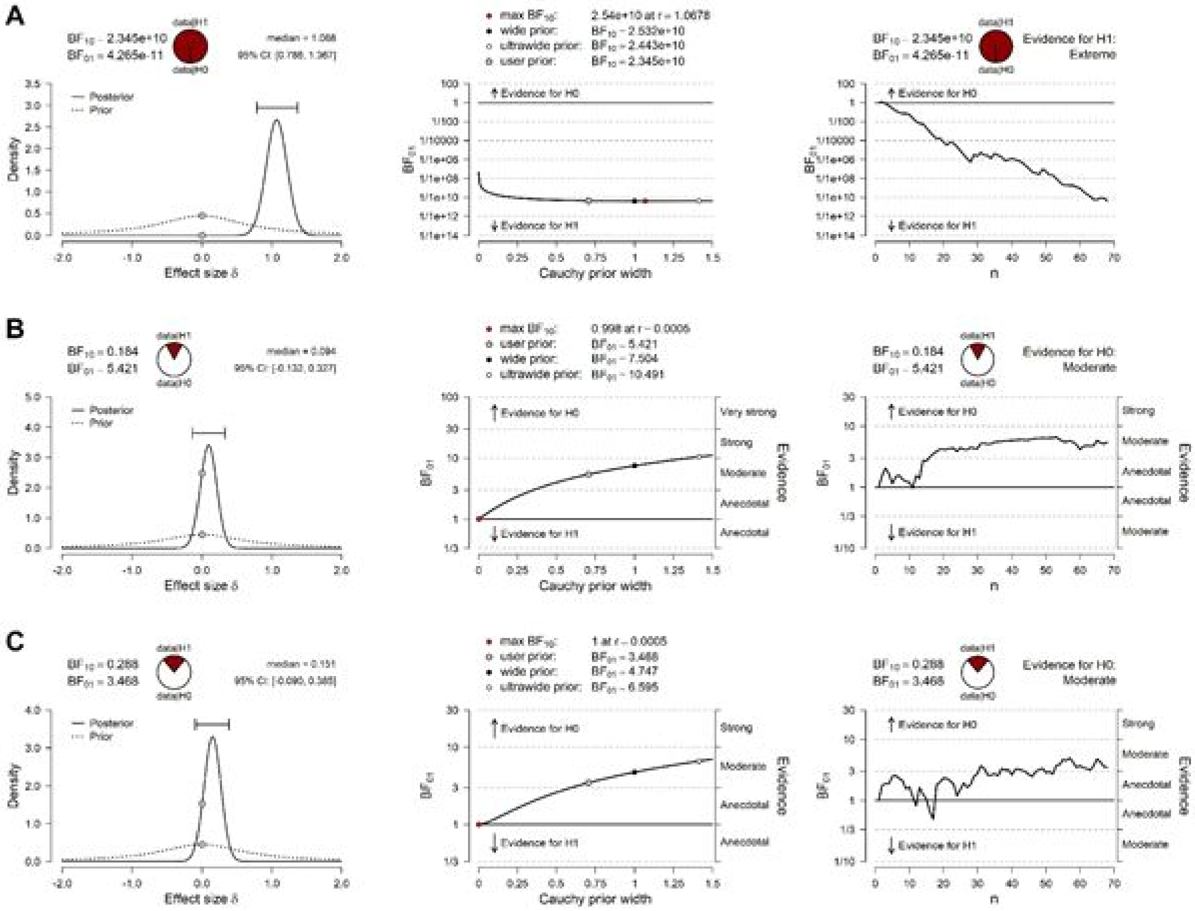


**Figure SM.5**. **Supporting plots for the EEG measures**. Panel A: Prior and posterior distributions, robustness checks, and sequential analysis for the *t*-test conducted on the N400 difference wave values. Panel B: Prior and posterior distributions, robustness checks, and sequential analysis for the *t*-test conducted on the LPC difference wave values. Panel C: Prior and posterior distributions, robustness checks, and sequential analysis for the *t*-test conducted on the hit-hit and hit-miss LPC difference wave values. Robustness checks show how the resulting Bayes factor changes across a wide range of priors. Sequential analysis depict the evidence for each of the null and alternative hypotheses, as sample size increases. Accompanying Bayes factors and the credible interval are also displayed.

### Frequentist Analyses

#### Behavioural Analyses

Six separate independent-samples *t*-tests were used to examine the impact of Val66Met genotype on the behavioural scores derived from the source memory task. For each test, we restricted the direction of testing, based on previous evidence that Val/Vals score higher on recognition tasks compared to Met allele carriers. For our measure of Discriminability, we found evidence that Val/Vals are better at discriminating old items from new items ($t(66) = -2.3$, $p = 0.01$, $d = -0.56$, 95%CI = $-\infty$; -0.15). Similarly we also find evidence for Val/Val individuals outperforming Met allele carriers on the item recognition task ($t(66) = -1.8$, $p = 0.04$, $d = -0.44$, 95%CI = $-\infty$; -0.33). However, there was no significant difference between the two groups for overall Hits ($t(66) = -1.5$, $p = 0.07$, $d = -0.36$, 95%CI = $-\infty$; 0.04), Correct Rejections ($t(66) = -1.6$, $p = 0.06$, $d = -0.39$, 95%CI = $-\infty$; 0.02), source recognition ($t(66) = -1.1$, $p = 0.15$, $d = -0.56$, 95%CI = $-\infty$; 0.15), or Response Bias ($t(66) = 0.29$, $p = 0.61$, $d = 0.07$, 95%CI = $-\infty$; 0.47).

#### N400 Component

In order to test whether we replicated the old/new effects previously described at the N400 time window, we ran a paired-samples *t*-test comparing the mean amplitude of ERPs to correctly identified old items to ERPs to correctly identified new items. We find correctly recognised old items to have more positive average amplitude across the N400 time window compared to correctly identified new items ($t(67) = 9.0$, $p = <0.001$, $d = 1.1$, 95%CI = 0.79; 1.39).

**Genetic impact on N400**. In order to examine whether there were any differences in the magnitude of the old/new effect for each of the Val66Met genotypes, we conducted an independent-samples *t*-test on mean difference amplitude for correctly identified old items and correctly identified new items, across the N400 time window. We restricted the direction of the *t*-test so that we could specifically examine whether Met allele carriers had smaller old/new effects compared to Val homozygotes. We find no genotype effect on the old/new effect for the N400 time window ($t(66) = -1.11$, $p = 0.14$, $d = -0.27$, 95%CI = $-\infty$; 0.13).

**Late Positive Component**

In order to test whether we replicated the old/new effects previously described at the LPC time window, we ran a paired-samples *t*-test comparing the mean amplitude of ERPs to correctly identified old items to ERPs to correctly identified new items. We find no difference between the average amplitude of ERPs in response to correctly recognised old items, compared to correctly identified new items ($t(67) = 0.83$, $p = 0.41$, $d = 0.1$, 95%CI = -1.39; 0.34).

**Genetic impact on LPC**. In order to examine whether there were any differences in the magnitude of the old/new effect for each of the Val66Met genotypes, we conducted an independent-samples *t*-test on mean difference amplitude for correctly identified old items and correctly identified new items, across the LPC time window. We restricted the direction of the *t*-test so that we could specifically examine whether Met allele carriers had smaller old/new effects compared to Val homozygotes, which we do confirm ($t(66) = -2.39$, $p = 0.01$, $d = -0.58$, 95%CI = $-\infty$; -0.17). Additionally, in this same time window, we were interested in testing whether there was a genotype influence on the average amplitude of ERPs that correspond to items that are correctly recognised as old with correct source judgment, and items that are correctly recognised as old with incorrect source judgment ($t(66) = -0.40$, $p = 0.35$, $d = -0.10$, 95%CI = $-\infty$; 0.31).
